# Tissue specific targeting of DNA nanodevices in a multicellular living organism

**DOI:** 10.1101/2021.03.05.434169

**Authors:** Kasturi Chakraborty, Sunaina Surana, Simona Martin, Jihad Aburas, Sandrine Moutel, Franck Perez, Sandhya P. Koushika, Paschalis Kratsios, Yamuna Krishnan

## Abstract

Nucleic acid nanodevices present great potential as agents for logic-based therapeutic intervention as well as in basic biology. Often, however, the disease targets that need corrective action are localized in specific organs and thus realizing the full potential of DNA nanodevices also requires ways to target them to specific cell-types *in vivo*. Here we show that by exploiting either endogenous or synthetic receptor-ligand interactions and by leveraging the biological barriers presented by the organism, we can target extraneously introduced DNA nanodevices to specific cell types in *C. elegans*, with sub-cellular precision. The amenability of DNA nanostructures to tissue-specific targeting *in vivo* significantly expands their utility in biomedical applications and discovery biology.

## Introduction

DNA has proven to be a versatile molecular scaffold to build an array of programmable synthetic nanoarchitectures due to its structural predictability(Seeman & Sleiman, 2017). The specificity of Watson-Crick base pairing, its tuneable affinity, the well-defined structural properties of the double helix and its modular nature make the DNA scaffold highly engineerable. A wide array of functional DNA-based nanodevices have been deployed *in vivo* both for quantitative chemical imaging and also as programmable carriers that deliver encapsulated cargo upon receipt of a molecular cue(Chakraborty, Leung, & Krishnan, 2017; Chakraborty, Veetil, Jaffrey, & Krishnan, 2016; Douglas, Bachelet, & Church, 2012; Jani, Zou, Veetil, & Krishnan, 2020; Krishnan & Bathe, 2012; Lee et al., 2012; J. Li et al., 2011; Modi et al., 2009; Narayanaswamy et al., 2019; Saha, Prakash, Halder, Chakraborty, & Krishnan, 2015; Sharma, Zaveri, Visweswariah, & Krishnan, 2014; Surana, Bhat, Koushika, & Krishnan, 2011; Thekkan et al., 2019; Veetil et al., 2017; Zhao et al., 2012). However, in most contexts, the site for payload action ideally needs to be confined to the desired organ or tissue. Thus, if DNA nanodevices can be targeted tissue-specifically in a live, multicellular organism, it would significantly expand their potential utility in biomedicine and fundamental biology.

Nature solves the problem of transporting poorly permeable molecules across membrane barriers, either releasing or enriching them tissue-specifically, by membrane trafficking. DNA nanostructures are particularly amenable to endosomal trafficking, which has in part, led to an array of synthetic DNA nanostructures being presently deployed in living systems(Bujold, Lacroix, & Sleiman, 2018; Chakraborty et al., 2016; Krishnan, Zou, & Jani, 2020). Endosomal trafficking is ubiquitous: it regulates the internalisation, sorting and intracellular transport of diverse cargo, and is pivotal to development, signalling and homeostasis. The significance and impact of endocytosis in health and disease is underscored by the observation that perturbations in endosomal trafficking are linked to multiple diseases, including neurodegeneration, cancer and cardiovascular disease and further, distinct cell types differ in their susceptibility to disease(Maxfield, 2014; Mellman & Yarden, 2013; Mukherjee, Ghosh, & Maxfield, 1997). By leveraging membrane trafficking as well as taking advantage of biological barriers present in the whole organism, we show that DNA nanodevices can be targeted tissue-specifically and with organelle-level precision in the nematode *Caenorhabditis elegans*.

Most studies that use DNA nanostructures as reporters or payload carriers describe their functionality primarily in cultured cells that express scavenger receptors for which DNA is the natural ligand(J. Li et al., 2011; Modi et al., 2009). *In vivo* studies in multicellular organisms have also exploited scavenger receptor-mediated endocytosis to target DNA architectures to phagocytic cells such as coelomocytes in *C. elegans(Surana et al., 2011)* or microglia in *D. rerio(Veetil et al., 2020)* (**Figure 1a**). However, since most cells and tissues do not endogenously express, scavenger receptors, targeting DNA nanostructures to such cell types *in vivo* remains challenging. In cultured cells, this problem is circumvented by exploiting naturally occurring receptor-ligand interactions. The natural ligand is chemically conjugated to a DNA nanodevice, which then binds its cognate receptor on the cell surface and gets internalized and transported within the cell along the trafficking route adopted by the receptor(Bhatia et al., 2016; Jani et al., 2020; Modi, Nizak, Surana, Halder, & Krishnan, 2013). Alternately, a synthetic receptor-ligand interaction has been used where the receptor is a sequence-specific, DNA-binding protein based on a single-chain variable fragment (scFv)(Modi et al., 2013). When the scFv is fused to a trafficking protein such as furin, and the chimera is expressed in cells, furin displays the synthetic scFv receptor on the cell surface. Thus, DNA nanodevices with an scFv-binding sequence engage the scFv domain of the chimera and get trafficked to specific organelles within the cell(Modi et al., 2013; Saminathan et al., 2020). Despite these *in cellulo* demonstrations, there is still no evidence that DNA nanostructures can be targeted to tissues lacking scavenger receptors in multicellular organisms.

**Figure 1:**
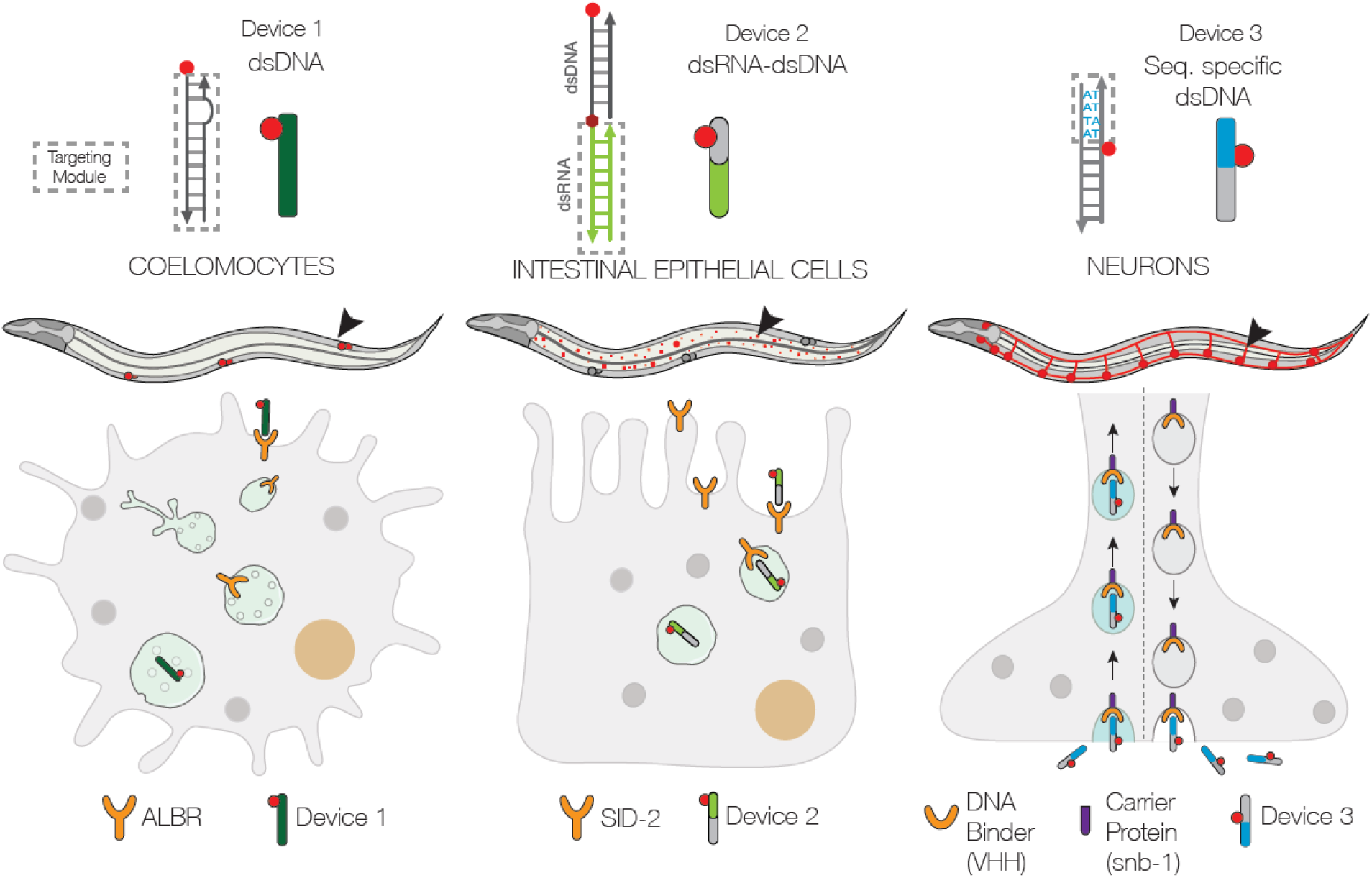
Schematic of strategies to target DNA devices to different cell types: (A) DNA nanodevices are intrinsically targeted to coelomocytes via the endogenously expressed scavenger receptors. (B) DNA nanodevices that display a dsRNA (green) domain are targeted to intestinal epithelial cells by engaging endogenously expressed SID-2 receptor. (C) DNA nanodevices are targeted selectively to those neurons that express a DNA-binding protein (V_H_H) fused to synaptobrevin (*snb-1*). The DNA nanodevice has a sequence (blue) recognized specifically by V_H_H.

In this study, we demonstrate that DNA nanodevices can be targeted tissue-specifically in the nematode *C. elegans* by leveraging both natural and synthetic receptors, on the surface of different cell types. We have focussed on two tissues where endosomal trafficking has been linked to critical physiological functions, namely, the intestine and the nervous system. In the first instance, we exploit the presence of endogenous SID-2 receptors on the intestine to target a DNA nanostructure along the endo-lysosomal pathway in intestinal epithelial cells (**Figure 1b**). In the second, we present a generalizable route to target DNA nanostructures to cells lacking either scavenger receptors or SID-2 receptors. Here, we use a synthetic receptor, namely, a newly identified, recombinant, single-domain antibody (9E) which tightly binds a specific 4-nt sequence of dsDNA. When 9E is fused to the synaptic vesicle protein synaptobrevin-1 (SNB-1) and selectively expressed in neurons, the SNB-1::9E chimera binds DNA nanodevices having the cognate 4-nt domain and localizes them in retrogradely-moving endosomes in neurons (**Figure 1c**).

By leveraging the *C. elegans* amenability to transgenesis, we demonstrate the molecular adaptability of this strategy. By expressing SNB-1::9E under promoters expressing in specific sets of neurons, we show that DNA nanodevices can be targeted to retrogradely-moving endosomes in specific neurons. We then demonstrate sub-cellular control over targeting afforded by this synthetic system. We show that DNA nanodevices can be displayed on the neuronal surface without invoking their entry into organelles. When 9E is genetically fused to the transmembrane odorant receptor ODR-2 and expressed under a neuron-specific promoter, it binds and positions a DNA nanodevice on the neuronal surface. The small size, stability and pH-insensitivity of 9E makes this two-component system immediately suitable for a multitude of DNA architectures bearing the cognate 4-nt sequence. Moreover, it offers a versatile route to target diverse proteins or track endosomal transport trajectories in different cell types *in vivo*, especially in model organisms amenable to transgenesis.

## Results and Discussion

### Targeting DNA nanodevices to intestinal epithelial cells

The intestine is one of the major organs of *C. elegans*, comprising a third of the total somatic mass with twenty large epithelial cells positioned with bilateral symmetry, forming a long tube around a lumen^(McGhee, 2007),^(Altun & Hall, 2009). Intestinal epithelial cells (IECs) contain prominent, birefringent gut granules that are known as lysosome-related organelles (LROs). Distinguished by lysosomal markers(Coburn & Gems, 2013; Dell’Angelica, Mullins, Caplan, & Bonifacino, 2000; Hermann et al., 2005), LROs play important roles in production and storage of melanin, immune defence and neurological function(Huizing, Helip-Wooley, Westbroek, Gunay-Aygun, & Gahl, 2008). We first sought to target a simple DNA nanodevice comprising a 38-bp double stranded DNA (dsDNA) bearing a 5’ Alexa 647 (A647), denoted D^38^, to label LROs in the intestinal epithelia (**Table 1**, **Supplementary Information**). We reasoned that the biological barrier between the intestinal lumen and the pseudocoelom would effectively preclude D^38^ from access to coelomocytes, as seen whenever DNA nanodevices are microinjected in the pseudocoelom^(Bhatia, Surana, Chakraborty, Koushika, & Krishnan, 2011; Surana et al., 2011)^.

**Table 1.**
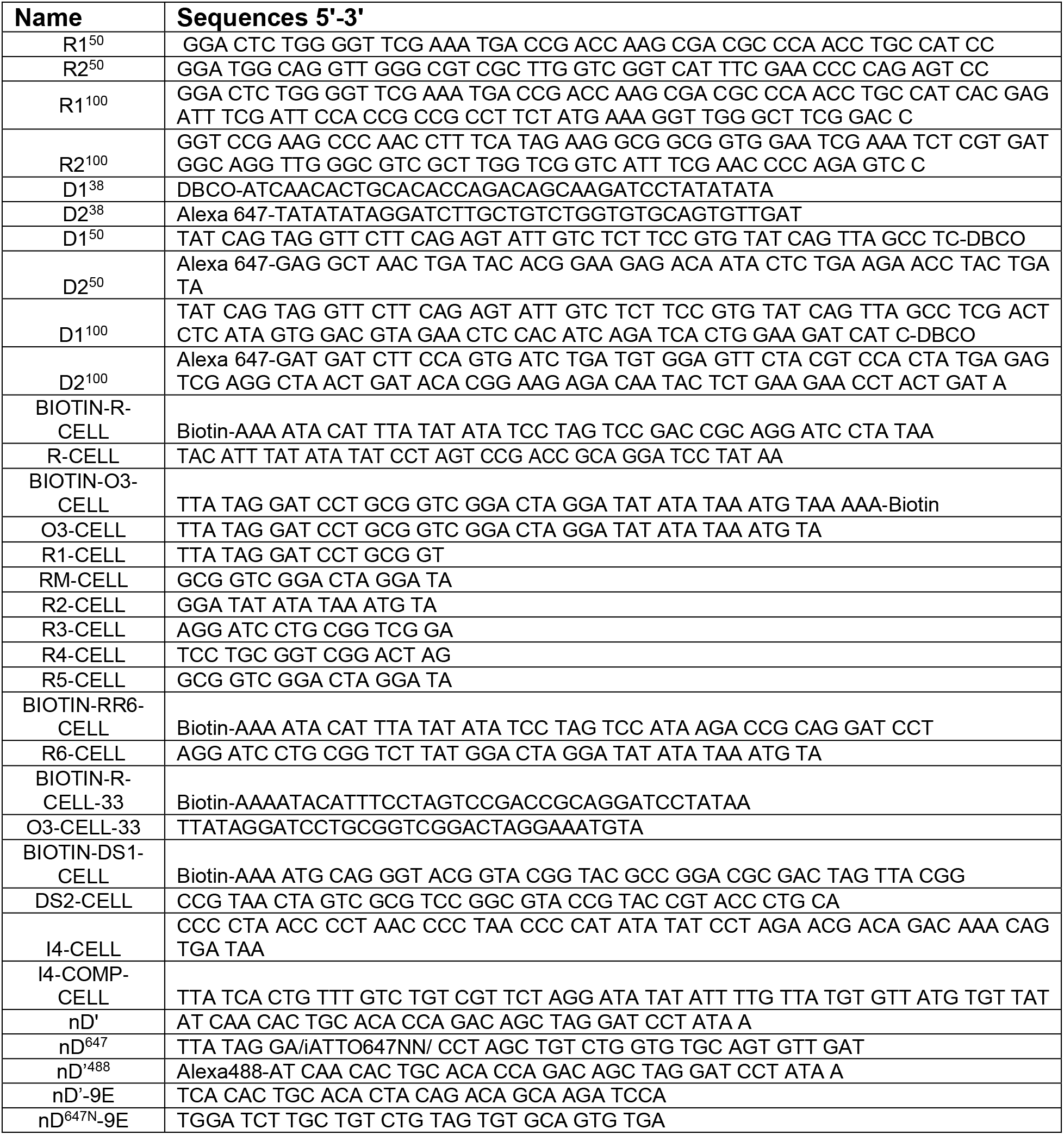
Sequences of oligonucleotides employed in this study.

We therefore utilized a liquid-feeding method to introduce 1 μM of D^38^ into the intestinal lumen of one-day adult worms (M&M). We leveraged the high autofluorescence of LROs at shorter wavelengths to provided regions of interest (ROIs) to evaluate the efficacy of D^38^ uptake. The mean intensity corresponding to D^38^ uptake, compared to control mock-fed worms, was very low, indicating low or no intrinsic targetability of DNA nanodevices to IECs. (**Figure 2a-b**). We therefore varied the size of the DNA nanodevice to test whether increasing the number of base pairs (bp) and/or negative charges improved targeting, the sequences of which are shown in **Table 1**, **Supplementary Information**. Using the same liquid-feeding method, we repeated uptake experiments with 50 or 100 bp long dsDNA (D^50^ or D^100^). Again, there was no enhancement in nanodevice targeting to IECs (**Figure 2a**). This revealed DNA nanostructures are not inherently targeted to cells in this tissue.

**Figure 2:**
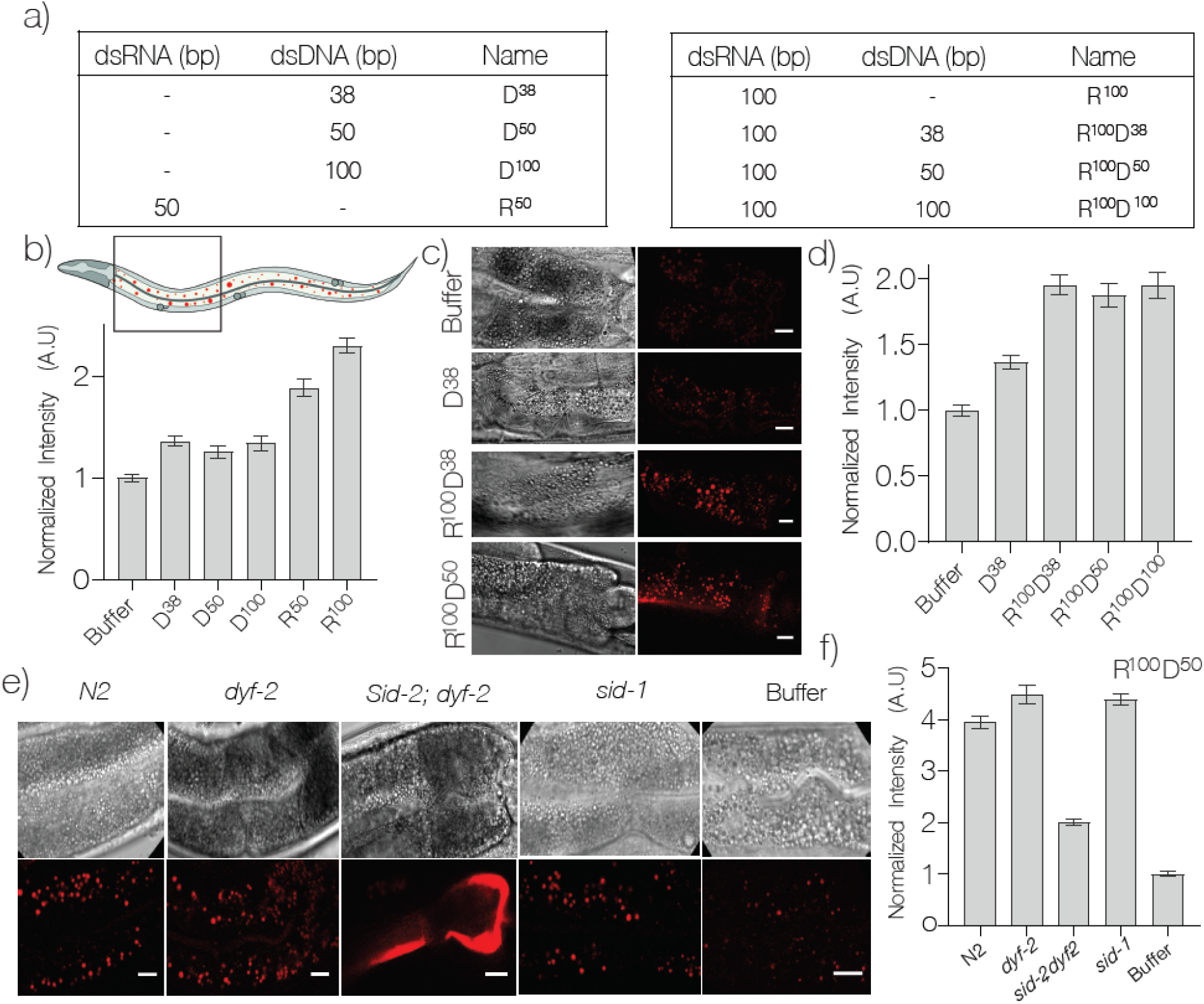
DNA nanodevices are targeted to intestinal epithelial cells. a) Table of the composition and length of the various Alexa-647 labeled DNA nanodevices used. B) Mean Alexa647 fluorescence intensity of corresponding to the uptake of the indicated nanodevice in *C. elegans* intestinal epithelial cells (IECs). c) Representative fluorescence and DIC images of IECs labelled with the indicated R^100^-conjugated nanodevice quantified in d). e) Representative fluorescence images of IECs labelled with nanodevice R^100^D^50^ in the indicated genetic background quantified in g). Data represented as mean + standard error of the mean of 10 worms, ~500 endosomes. The four IECs closest to the pharynx were considered for quantification. Scale bar = 10 μm dsDNA scaffold had no significant effect on uptake efficiency (**Figure 2d**-**e)**. Thus, it is possible to target DNA nanodevices to intestinal epithelial cells using a DNA-RNA chimera.

We then tested whether we could co-opt the RNA interference (RNAi) pathway in order to target DNA-based probes to intestinal epithelia. Previous studies have shown that worms are capable of endocytosing dsRNA localized in the acidic intestinal lumen, and that this is effected by the membrane protein SID-2(Hunter et al., 2006; McEwan, Weisman, & Hunter, 2012; William M Winston, Sutherlin, Wright, Feinberg, & Hunter, 2007). We first tested the uptake of dsRNAs either 50 bp or 100 bp long, denoted R^50^ or R^100^ respectively, based on previous work suggesting this as the minimum length needed for effective internalisation by SID-2 (**Figure 2a**)(McEwan et al., 2012)All dsRNAs carried a fluorescent Alexa647 label at their 5’ termini. Interestingly, when worms were liquid-fed either R^100^ or R^50^ we observed that uptake efficiency by intestinal epithelia increased ~2-fold compared to their DNA analogues (**Figure 2b**).

We therefore sought to test whether covalently attaching R^100^ or R^50^ to a DNA nanodevice would promote nanodevice entry into the intestinal epithelia via the SID-2 receptor, and potentially label LROs. Using click chemistry, we conjugated R^100^ to DNA nanodevices D^38^, D^50^ or D^100^ bearing Alexa647 labels to give chimeric RNA-DNA nanodevices denoted R^100^D^38^, R^100^D^50^ and R^100^D^100^ respectively (**Figure 2c**-**d**, **Supplementary Fig 1**)(Jewett, Sletten, & Bertozzi, 2010). When nematodes were liquid-fed the above chimeras and then imaged, we found that overall, conjugating R^100^ to DNA nanodevices improved uptake of the latter by intestinal epithelial cells. However, interestingly, we observed that the length of the

Closer scrutiny revealed that R^100^D^n^ nanodevices internalized by intestinal epithelia localized to lysosome related organelles (LROs), readily identified by their high autofluorescence at low wavelengths(Soukas, Carr, & Ruvkun, 2013), as well as colocalization with LAMP-1::GFP and GLO::1-GFP (**Supplementary Fig 1)(Schroeder et al., 2007; Soukas et al., 2013)**. In order to confirm the identity of the receptor responsible for the uptake of the chimeric nanodevices into LROs, we repeated uptake assays in worms carrying mutant alleles for *sid-1* and *sid-2*, two critical players of the RNAi pathway. Homozygous animals for the *sid-1(qt9)* null allele are systemically resistant to RNAi(W M Winston, Molodowitch, & Hunter, 2002). The s*id-2* locus is embedded within an intron of another gene (*dyf-2*) and the only available strong loss-of-function allele (*gk505*) for *sid-2* contains a 403 bp deletion, which removes the first *sid-2* exon and the twelfth *dyf-2* exon. As a control for the *sid-2&dyf-2(gk505)* strain, we used animals carrying the *dyf-2(gk678)* mutant allele that selectively disrupts the *dyf-2* gene alone. Wild-type (N2 strain), *dyf-2, sid-1, and sid-2&dyf-2* worms were tested for their ability to uptake R^100^D^50^ as previously described (**Figure 2d**-**e)**. We found that uptake was reduced in *sid-2&dyf-2* mutants, but not in *sid-1(qt9)*, *dyf-2(gk678)* or wild-type worms. Our results are consistent with SID-2 being localized on the extracellular surface, acting as a receptor that binds and endocytoses dsRNA, while SID-1 is resident in endosomes, where it binds dsRNA cargo and transports it into the cytosol(Jose & Hunter, 2007; W. Li, Koutmou, Leahy, & Li, 2015; McEwan et al., 2012). This implicates SID-2 as the receptor for nanodevice internalisation and its subsequent trafficking to LROs in intestinal epithelial cells.

### Development of a DNA binding, recombinant camelid antibody

*C. elegans* has proven to be a powerful model organism to study vesicular trafficking in neurons, given its transparent body and simple nervous system(Hulme & Whitesides, 2011; Teschendorf & Link, 2009). However, neurons do not express either scavenger receptors or SID proteins, complicating the targeting of DNA nanodevices to these cell types. We therefore developed a DNA-binding recombinant, camelid antibody (V_H_H) as a synthetic receptor for DNA nanodevices. This offers the advantage of tagging endogenously expressed proteins in any cell of choice rather than ectopically expressing canonical DNA-binding proteins that could have unknown biological effects. The small size of camelid antibodies, the ability to select them by display technologies and tune their affinity, stability, and expression by molecular evolution have led to a plethora of *in vivo* applications(Antibodies, Phage, & Technology, 1994). Camelid antibodies lack a light chain altogether and the heavy chain domain itself is sufficient to form the antigen binding pocket(Hamers-Casterman et al., 1993), leading to their utility in therapeutics as well as cell biology(Harmsen & De Haard, 2007). They robustly maintain their native conformations due to increased hydrophilicity and single domain nature and are much more resistant to thermal and chemical denaturation(Dumoulin et al., 2002; Ewert, Cambillau, Conrath, & Plückthun, 2002; Pérez et al., 2001; van der Linden et al., 1999).

We screened a recombinant llama antibody library to isolate high affinity candidates that bind a specific DNA duplex and characterized the binding *in vitro*^5^. After a phage display screen, we studied 160 potential binders and selected one of the highest affinity binders for further evaluation for its affinity and specificity of binding. This binder, 9E, was further tested for its expression, pH dependence and binding affinity. To date, V_H_H recombinant antibodies have been obtained using phage display against many classes of molecules, but not against nucleic acids.

We therefore developed an assay to screen recombinant camelid antibodies by phage display against a 41 bp dsDNA to obtain high-affinity, sequence-specific DNA binding antibodies (**Materials and Methods**). The 41 bp dsDNA target was immobilized on streptavidin coated magnetic beads via a 5′ biotinylated terminus and presented as the epitope to the V_H_H library (**Figure 3a**). Following standard procedure with the variations described above, three rounds of progressive selection and amplification enriched for putative dsDNA binders. Next, 160 clones, were randomly selected, grown in 96 well plates and the corresponding phage bound V_H_H antibodies were expressed and screened for dsDNA binders. Nearly 70% displayed DNA binding properties, of which 40 bound DNA regardless of whether it was double- or single-stranded. Interestingly, 42 were found to show sequence specificity for the dsDNA target and showed minimal binding to ssDNA of the same sequence or dsDNA with a different sequence. These 42 clones were taken forwards for further analysis (**Supplementary Figure 2**).

**Figure 3:**
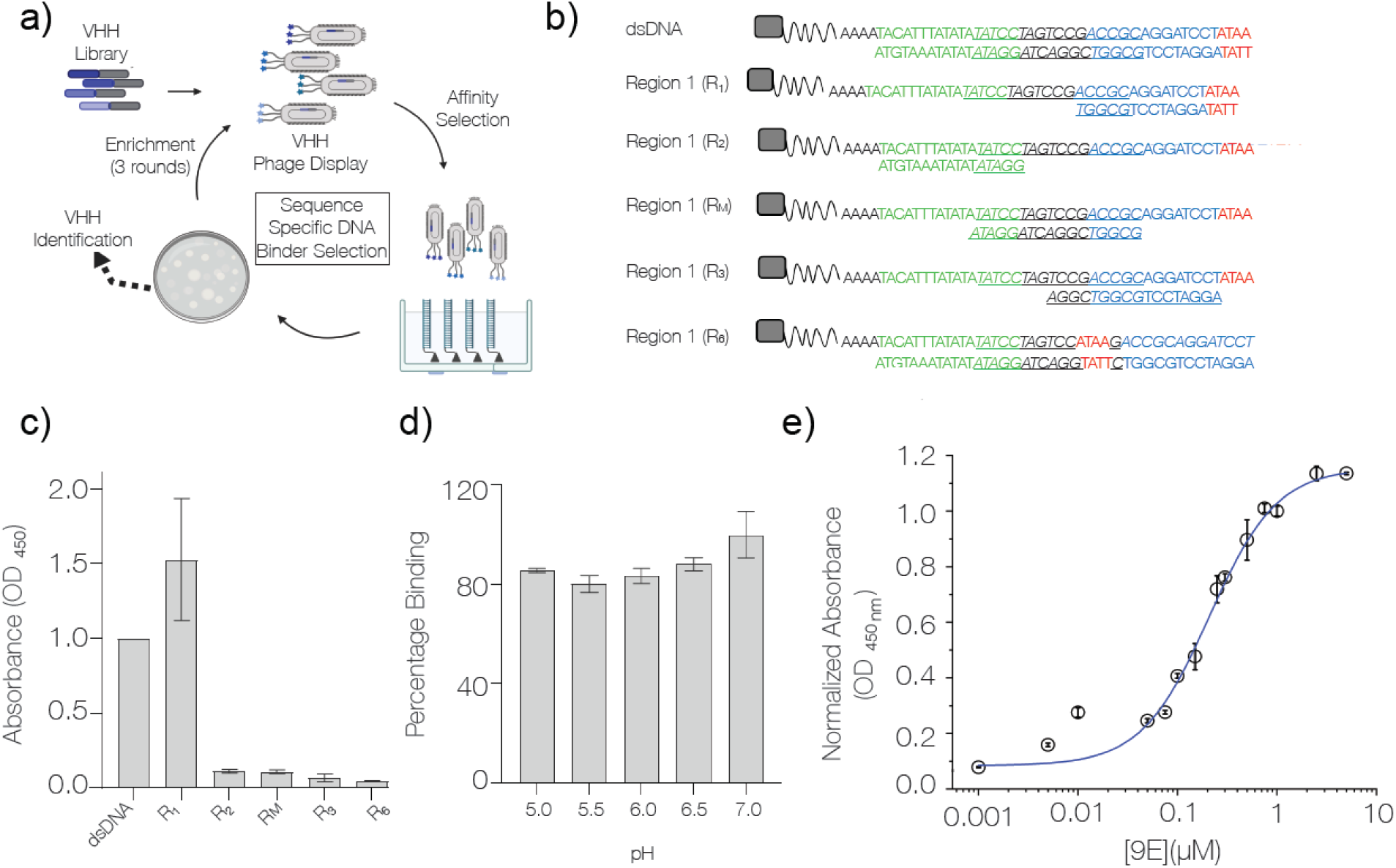
Identification and characterization of a sequence-specific DNA-binding recombinant antibody 9E. a) Schematic of phage display screen to identify DNA binders using a camelid antibody (VHH) library b) Sequence of the various dsDNA epitopes used to pinpoint the dsDNA sequence bound by the recombinant antibody, 9E. The tested regions are R_1_ (blue and red), R_2_ (green), an overlapping region R_M_ (italicized and underlined), R_3_ (region R_1_ frame-shifted by a window of 4 nts) and R_6_ (4-nt motif in red, present in the middle of the DNA duplex). Biotin (grey square) was incorporated to immobilize the DNA epitopes on streptavidin-coated magnetic beads. c) Binding efficiencies of the recombinant antibody 9E with the indicated dsDNA constructs as determined by ELISA. d) Effect of pH on binding efficiency of 9E with dsDNA as determined by ELISA. e) The relative binding constant of 9E and dsDNA epitope in solution. Serial dilutions of the purified protein were added to a fixed amount of immobilized dsDNA (25 pmoles). All experiments were performed in triplicate and the data is represented as mean ± s.e.m.

To screen for the minimal dsDNA motif recognized by each of the selected V_H_H antibody clones, ELISA was performed against a set of immobilized dsDNAs. These corresponded to the full 41-mer dsDNA target and three shorter dsDNA regions R_1_, R_2_ and R_M_ on the target (**Figure 3b**). Region R_1_ corresponded to a 17 bp region at the 3′ terminus of the biotinylated ssDNA oligonucleotide (nucleotides in red and blue font), R_2_ corresponded to a 17 bp region on the 5’ end of the oligonucleotide (nucleotides in green font), while R_M_ corresponded to a 17 bp overlapping the 5′ end of R_1_ and the 3′ of R_2_ shown in italics. Of the 42-dsDNA binding V_H_H antibodies tested, nearly 80% specifically bound region R_1_, albeit with varying affinities (**Supplementary Figure 3**).

To pinpoint the epitope on the 17-bp region R_1_, partial duplexes were tested covering this region with a sliding window of 4 bp from the 3’ end of the biotinylated oligonucleotide to form dsDNAs R_3_ and R_6_ (**Figure 3b**). This strategy ensured that the epitope could be narrowed down to at least 4 bp. ELISAs revealed that each of the 42 V_H_H antibodies required only the first 4 bp at the 3’ end of region R_1_ (ATAA, nucleotides in red) for binding (**Figure 3c**). When this 4 bp sequence is present not at the terminus, but in the middle of the dsDNA duplex, as in R_6_, it was no longer a ligand for any of the 42 V_H_H antibodies. This clearly demonstrated that the minimal binding epitope for all 42 V_H_H antibodies isolated was a terminal d(ATAA) motif (**Supplementary Figure 3**). Furthermore, since endocytic events are accompanied by lumenal pH changes, it is crucial to test whether binding between the V_H_H antibody to its cognate DNA epitope is pH-independent. From a pH-dependent ELISA screen we identified that the V_H_H 9E was disrupted the least of all by acidification, demonstrating ~82% binding even at pH 5.0. (**Figure 3d, Supplementary Figure 4**). Finally, based on ELISA, we estimated that the relative binding affinity of V_H_H 9E to its target epitope was ~200 nM (**Figure 3e, Materials and Methods**). Cumulatively, our findings show that the recombinant camelid antibody V_H_H 9E binds with high specificity to a 3’ terminal, d(ATAA) region (**Supplementary Figure 5**).

### Pan-neuronal expression of the SNB-1::9E chimera in C. elegans

Next, we tested whether DNA nanodevices could be targeted to neuronal endosomes by genetically fusing 9E, the DNA-binding V_H_H antibody identified above, with a protein such as synaptobrevin (SNB-1), that is expressed in neurons and undergoes endosomal trafficking. Our choice of synaptobrevin as a carrier protein was guided by it being an integral membrane protein present in multiple copies in synaptic vesicles(Dittman & Ryan, 2009). It has a low molecular weight (~16 kDa) and its N-terminal cytoplasmic domain interacts with target membrane SNARE (t-SNARE) proteins to facilitate neurotransmitter exocytosis, while its C-terminus has a hydrophobic patch that anchors it to vesicular membranes. The C-terminus ends in a short tail which extends into the vesicular lumen and is exposed to the synaptic cleft during exocytosis(Hanson, Heuser, & Jahn, 1997; Südhof, 1995). Upon their fusion with the plasma membrane at the synaptic cleft, synaptic vesicles undergo recycling via the endosomal pathway(Rizzoli, 2014). Furthermore, fusing GFP (~27 kDa) to the C-terminus of SNB-1(Murthy, Bhat, & Koushika, 2011; Nonet, 1999), does not perturb SNB-1 localization or function(Nonet, 1999) (**Figure 1c**).

Therefore, we reasoned that neuronal expression of an SNB-1::9E fusion protein would lead to the 15 kDa V_H_H domain being displayed in the synaptic cleft upon neurotransmitter release. Thus, if DNA nanodevices bearing a terminal d(ATAA) motif were present in the pseudocoelom, they could bind the V_H_H domain (blue) of the chimera at the synaptic cleft and get trafficked retrogradely along the endosomal pathway (**Scheme 1**). We therefore generated transgenic *C. elegans* animals, denoted *Psnb-1:snb-1::9E* worms, that express the SNB-1::9E chimera in all neurons under the control of the *snb-1* promoter (p*snb-1*)(Stefanakis, Carrera, & Hobert, 2015), including an *unc-54* 3’ UTR for efficient translation (**Figure 4a-b, Supplementary Figure 6**). Into these worms we injected the DNA nanodevice D^38^ (1 μM) modified to display a terminal d(ATAA) motif and an Atto 647N fluorophore, now denoted nD^A647^, into the pseudocoelom and imaged the worms after 30 min. We observed several punctate structures, linearly arranged along the worm’s body strongly resembling neuronal synapses along the *C. elegans* dorsal and ventral nerve cords (**Figure 4b**).

**Figure 4:**
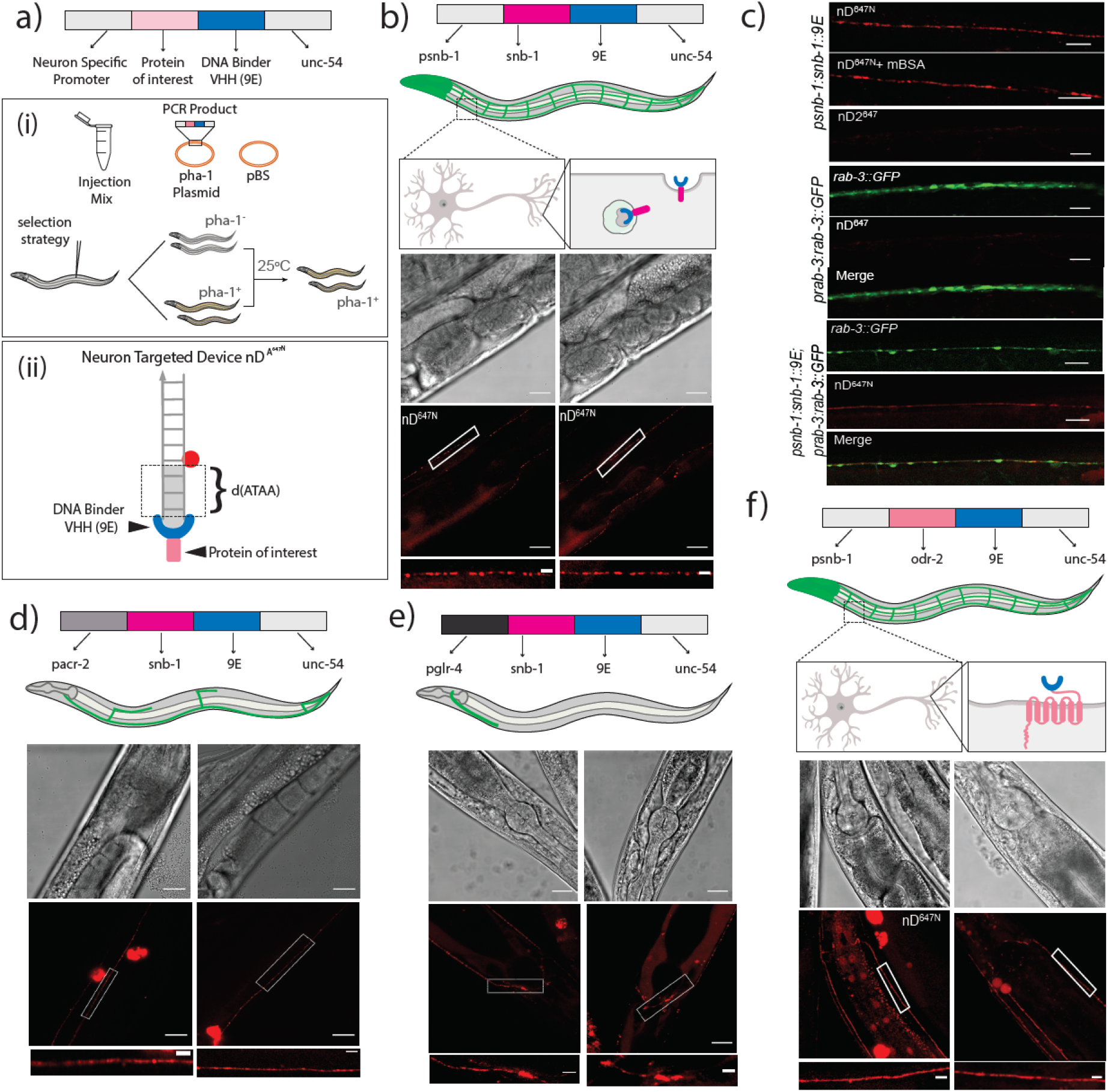
Targeting DNA nanodevices to neurons. a) Schematic of the constructs used to make transgenics i) Strategy to select transgenics based on *pha-1+* worms ii) Schematic of neuron targetable DNA nanodevice nD^A647^ bound to its synthetic receptor, 9E, fused to the protein of interest. b) Schematic of nanodevice uptake into neurons of *psnb-1:snb-1::9E* worms. Brightfield and fluorescence images of *C. elegans* neurons labeled with nD^A647^. c) Representative fluorescence images of i) neurons in *psnb1:snb-1::9E* worms injected with nD^A647^ in the presence or absence of mBSA and nD2^A647^ lacking the 3’ terminal d(ATAA) ii) neurons in *prab-3::gfp::rab-3* and iii) *prab-3::gfp::rab-3*; *psnb1:snb-1::9E* expressing *C. elegans* injected with nD^A647^. nD^A647^ can be targeted to neuronal subsets. d) nD^A647^ labels cholinergic neurons in *pacr-2:snb-1::9E* and glutamergic neurons in e) *pglr-4:snb-1::9E* expressing *C. elegans*. nD^A647^ can be targeted via the labeling cassette to other proteins cell-specifically. f) Images of neurons in *psnb1::odr-2::9E* expressing animals labeled with nD^A647^.. In all images, the white boxedregion is shown in the zoomed image below. Scale: 20 μm; Inset: 5 μm.

To test whether the nanodevices were in fact labelling neurons and synapses, we performed a series of colocalization experiments using transgenic *C. elegans* animals carrying different neuronal and synaptic markers. The strain *otIs355 [rab-3p::2xNLS::tagRFP]* expresses nuclear RFP pan-neuronally, whereas *otIs45 [unc-119p::GFP]* expresses cytosolic GFP pan-neuronally (**Supplementary Figure 7)***(Altun-Gultekin et al., 2001; Nguyen et al., 2016)*. The *jsIs682 [rab-3p::gfp::rab-3]* strain expresses GFP::RAB-3 in most neurons, which is localized primarily to synaptic regions (**Supplementary Figure 7)(Mahoney et al., 2006)**. Hermaphrodites carrying the *Psnb-1:SNB-1::9E* transgene were crossed with each of the aforementioned reporter strains (**Supplementary Figure 6)**. When 500 nM nD^A647^ was injected into *jsIs682 [rab-3p::GFP::RAB-3]; Psnb-1:SNB-1::9E* worms, we found that nD^A647N^ labelling coincides with RAB-3:GFP, confirming synaptic labelling. Interestingly, nD^A647N^ containing puncta do not exclusively colocalize with RAB-3:GFP positive puncta within the same axon. This suggests that in addition to the synaptic regions of neurons, nD^A647N^ might be labelling retrogradely trafficking endosomal compartments or synaptic vesicles that are being recycled. Similar colocalization experiments of nD^A647^ with other neuronal markers such as *otIs355 [rab-3p::2xNLS::tagRFP]* and *otIs45 [unc-119p::GFP]* further confirmed that nD^A647N^ indeed labeled neurons (**Supplementary Figure 7**).

Normally when DNA nanodevices are introduced into the pseudocoelom, they are taken up by scavenger receptors present on coelomocytes(Chakraborty et al., 2017; Dan, Veetil, Chakraborty, & Krishnan, 2019; Narayanaswamy et al., 2019; Surana et al., 2011; Veetil et al., 2017). A key characteristic of nanodevice uptake via this pathway is that uptake is easily abolished in the presence of 10 equivalent molar excess of maleylated BSA, as the latter competes for scavenger receptors due to its anionic nature(Haberland & Fogelman, 1985). Importantly, even in the presence of 10 equivalents excess of maleylated BSA, neuronal labelling of *psnb-1:snb-1::9E* worms with nD^A647^ was not affected (**Figure 4c-i**). This indicates that nD^A647^ uptake into neurons does not occur via scavenger receptors. When the terminal d(ATAA) motif in nD^A647^ was removed to give a DNA nanodevice denoted nD2^A647^, we observed no such neuronal labelling (**Figure 4c-ii).**These results show that DNA nanodevices can be targeted to neurons in worms only if the former incorporate a d(ATAA) motif, and if the latter express the SNB-1::9E chimera in neurons.

### Targeting DNA nanodevices with cellular and subcellular precision

To assess the precision over targeting using our method, we sought to localize DNA nanodevices in endosomes of specific neuron types. We therefore expressed the SNB-1::9E chimera under two different neuronal promoters. We focused on the cholinergic subset of motor neurons since acetylcholine is the most broadly used neurotransmitter in the nematode nervous system(Pereira et al., 2015). Cholinergic motor neurons in the ventral nerve cord are divided into six classes: DA, DB, AS, VA, VB, and VC classes(Kerk, Kratsios, Hart, Mourao, & Hobert, 2017). Since the acetylcholine receptor subunit ACR-2 is expressed in four of the six classes of cholinergic motor neurons (DA, DB, VA, VB), we used the ACR-2 promoter region to target these subsets of neurons(Kratsios, Stolfi, Levine, & Hobert, 2011). Transgenic *Pacr-2::snb-1::9E* worms were generated using the previously described *pha*-*1* strategy (**Figure 4a**)(Granato, Schnabel, & Schnabel, 1994). When nD^A647^ (1 μM) was introduced into the pseudocoelom of these nematodes, we observed that it localized in several, linearly arranged punctate structures resembling synapses and/or neuronal endosomes, around the mid-section of the worm body corresponding to the axons of the VA and VB neurons of the ventral nerve cord (**Figure 4d**). The DA and DB neurons extend their axons to the dorsal nerve cord and form synapse with dorsal muscle. However, we did not observe any labelling with nD^A647^ in the dorsal nerve cord of *Pacr-2::SNB-1::9E* worms. Thus, our strategy selectively labels VA and VB neurons over DA and DB classes of neurons. Such selective labelling could arise from the differential cell surface levels of SNB-1::9E in VA and VB neurons, or greater accessibility of their synapses to cargo in the pseudocoelom, or both.

Next, we targeted head neurons and a distinct set of cholinergic motor neurons, namely the SAB class, located proximal to the pharynx that innervates the head muscles. In order to target nD^A647^ to these neurons, we chose the promoter of the glutamate receptor family protein GLR-4(Feng et al., 2006), since this protein is known to be expressed in the head neurons and the SAB, VB and DA9 neurons(Brockie, Madsen, Zheng, Mellem, & Maricq, 2001; “glr-4 (gene) - WormBase : Nematode Information Resource,” n.d.; Hills, Brockie, & Maricq, 2004; Kratsios et al., 2015; Rothaug et al., 2014). When nD^A647N^ (1 μM) was introduced into the pseudocoelom of these nematodes, we observed that it localized in punctate structures arranged around the pharynx, corresponding to the putative synapses of head neurons expressing *glr-4*, and ventral neuromuscular synapses belonging to the SAB and VB neurons (**Figure 4e**). Again, just as in the previous case of the DA and DB neurons, the DA9 neurons at the tail of the worm did not show labelling, revealing that this method led to the preferential labelling of SAB and VB neurons. Taken together, our results show that DNA nanodevices can be targeted to synapses and/or retrogradely trafficking endosomes of specific neuron types expressing SNB-1::9E.

We then tested whether our method provides sub-cellular level precision in terms of targeting individual neurons. Therefore, we tested whether we could target a DNA nanodevice to neurons, and yet not allow neuronal entry, thereby keeping the DNA nanodevice immobilized on the neuronal surface. We therefore genetically fused 9E to the ODR-2 protein and expressed it under the pan-neuronal p*snb-1* promoter (**Figure 4f**). ODR-2 is a GPI-anchored membrane-associated signalling protein that plays important roles in nematode chemotaxis(Chou, Bargmann, & Sengupta, 2001). ODR-2 is highly expressed in sensory neurons, motor neurons and interneurons, and is particularly enriched in axons(Gottschalk & Schafer, 2006). Using the *pha-1* selection strategy described earlier, we obtained transgenics stably expressing the ODR-2::9E chimera (**Figure 4a**). When these transgenics were injected with nD^A647N^ (1 μM), we observed that in all worms, the DNA nanodevice labels neurons corresponding to the ventral and the dorsal nerve cords (**Figure 4f**). In these transgenics, a near-uniform pattern of plasma membrane labelling was observed that is reminiscent of ODR-2 staining(Chou et al., 2001), and is in stark contrast to the punctate labelling that was observed in p*snb-1::snb-1::9E* worms (**Figure 4b**). Taken together, our experiments show that the synthetic receptor-ligand strategy shown here is generalizable to cell-surface membrane proteins and can be used to label specific membrane domains and/or endosomal compartments across neurons *in vivo*. Additionally, this design is applicable to a range of functional DNA nanostructures provided they harbour a d(ATAA) tag, opening up new avenues for the cell-specific application of DNA nanodevices in biological systems.

## Conclusion

Endocytosis and endosome trafficking are complex processes, regulating many critical cellular functions and therefore play pivotal roles in tissue-specific physiology and pathology. Our knowledge of subcellular dynamics and organization of endocytic and recycling pathways stems largely from investigations in cultured cellular systems. However, *in vivo*, endocytosis in single cells and their functional outcomes are sculpted by molecular interactions with their neighbouring cells, occurring in the presence of multiple biochemical cues originating from other tissues. Thus, the ability to study endocytic pathways *in vivo*, at the single cell level and in their native context, would illuminate our understanding of this core cellular process at the whole organism level.

There is now a rich repertoire of investigative tools based on DNA nanodevices that either probe or manipulate endosomal trafficking pathways(Krishnan et al., 2020). However, in order to tap the full potential of DNA-based nanodevices *in vivo*, it is essential to develop strategies for precise targeting to specific tissues. Here, we leverage both endogenous as well as synthetic nucleic acid-binding receptors to specifically target simple DNA nanodevices to two kinds of tissues: intestinal epithelial cells and neurons in *C. elegans*.

Our results show that, unlike what has been observed for coelomocytes(Chakraborty et al., 2017; Dan et al., 2019; Narayanaswamy et al., 2019; Surana et al., 2011; Veetil et al., 2017), simply introducing DNA nanodevices into the worm intestine and circumventing the relevant biological barrier, does not lead to nanodevice targeting to intestinal epithelial cells. This is due to the lack of an appropriate intestinal epithelial cell surface receptor for DNA. However, by exploiting the endogenous dsRNA receptor SID-2 in intestinal epithelial cells, we could target DNA nanodevices to these cells. We achieved this by conjugating a dsRNA domain to DNA nanodevices, where the dsRNA domain engaged the SID-2 receptor such that the DNA nanodevice underwent receptor-mediated endocytosis and was subsequently trafficked to lysosome related organelles (LRO). Furthermore, since mutations in the *sid-2* gene, but not *sid-1*, abrogated DNA nanodevice uptake, we could pinpoint that nanodevice targeting occurs via the endogenous SID-2 receptor. Our results show that DNA nanodevices can be targeted tissue-specifically by exploiting relevant endogenously expressed cell-surface receptors and displaying the cognate ligand on the DNA nanodevice. Should new knowledge of cell type-specific nucleic acid receptors, this strategy could be used to expand the targeting of DNA nanodevices to other classes of cells.

In the absence of such knowledge, we have developed a synthetic receptor-ligand strategy to target DNA nanodevices in a generalisable way so that it can be adapted to a variety of cellular and molecular contexts *in vivo*. We therefore identified a sequence-specific, DNA-binding recombinant camelid antibody, denoted 9E, by phage display. 9E binds a specific 4-nt sequence, namely d(ATAA) when it is present at the 3’ end of the DNA nanodevice with high affinity (~200 nM), irrespective of the environmental pH.

By fusing 9E to proteins that are expressed on the target cell under appropriate tissue-specific promoters, we can target DNA nanodevices with cellular and sub-cellular level precision. For example, since SNB-1 is ubiquitously expressed in all neurons, we expressed a SNB-1::9E fusion protein under the pan-neuronal promoter p*snb-1* and targeted a DNA nanodevice displaying the 3’ terminal d(ATAA) sequence, specifically to neurons. Interestingly, nanodevices labelled endosomes not only at the synapses but all along the axon within the relevant neurons. Neuronal targeting required both, SNB-1 fused to the 9E antibody and the DNA nanodevice to have the 3’ d(ATAA) sequence. This established that cell-specific targeting was contingent upon the interaction between the synthetic receptor and its cognate DNA epitope. DNA nanodevices could be targeted to specific sub-sets of neurons by expressing the SNB-1::9E chimera under promoters of genes expressed in specific neuronal classes. Thus, when SNB-1::9E was expressed under p*acr-2* or p*glr-4* promoters(Feng et al., 2006; Kratsios et al., 2015, 2011), DNA nanodevices were internalized by cholinergic ventral cord neurons or SAB head motor neurons respectively.

We showed that targeting of DNA nanodevices could be achieved with sub-cellular level precision using this strategy. Fusing 9E to the transmembrane odorant receptor, ODR-2, which does not undergo significant endocytosis(Chou et al., 2001), enabled the immobilization of a DNA nanodevice at the neuronal surface, rather than being endocytosed into intracellular vesicles. This suggests that DNA nanodevices may be localized with subcellular precision using a synthetic receptor and cognate ligand strategy *in vivo*.

To interrogate organelle dynamics and function *in vivo*, the development of robust technologies with molecular specificity that enables access to discrete cell types is critical. The modular system described here has the capability to interface with and harness, the power of nucleic acid nanotechnology to probe and program specific tissues *in vivo*. These targeting strategies open up an array of possible applications where DNA nanodevices can be readily applied in the multicellular context, and positions DNA nanodevices to deliver key insights into complex biological phenomena.

## Acknowledgements

We thank the Integrated Light Microscopy facility at the University of Chicago and the Caenorhabditis Genetic Center (CGC) funded by NIH Office of Research Infrastructure Programs (P40 OD010440) for strains. This work was supported by the University of Chicago Women’s Board; FA9550-19-0003 from AFOSR (YK), NIH grants 1R01NS112139-01A1 (YK), the Ono Pharma Foundation Breakthrough Science Award (YK) and a Whitehall Foundation Grant 2017-12-50 (PK).

## Author Contribution

SS, KC, SPK, PK and YK designed the project. SS, SM, FP and YK developed and characterized the recombinant 9E. KC and SM made constructs for transgenic development. KC, JA and PK developed all new transgenics. KC developed all the DNA nanodevices, performed biochemical assays, worm imaging and analysis. KC and YK analyzed the data. KC, SS and YK wrote the paper. All authors discussed the results and gave inputs on the manuscript.

## Competing interests

The authors declare no competing interests.

## Data availability

The data that support the plots within this paper and other finding of this study are available from the corresponding author upon reasonable request.

## Supplementary Information

### Oligonucleotides

All fluorescently labeled DNA oligonucleotides were HPLC-purified and obtained from IBA-GmBh (Germany) and IDT (Coralville, IA, USA). Unlabeled DNA oligonucleotides were purchased from IDT (Coralville, IA, USA). RNA was transcribed *in vitro* using a MEGAscript^®^ T7 Kit (Invitrogen, USA). geneBlock fragements were obtained from IDT (Coralville, IA, USA)

Sequences of the various oligonucleotides used are listed below in Table 1.

### Preparation of oligonucleotide samples

All oligonucleotides were ethanol precipitated, dissolved in Milli-Q water, aliquoted as a 100 μM stock and stored at −20°C. Concentration of each oligonucleotide was measured using UV absorbance at 260 nm.

An intestinal epithelial cell targeting probe consist of an azido labelled RNA duplex conjugated to a DBCO containing fluorophore or DNA duplex via copper free click chemistry^1–3^. A PCR fragment of 100 bp or 50 bp template with a T7 promoter site obtained from a plasmid DNA was used as a template for *in vitro* transcription using protocols suggested by the manufacturer (MEGAscript^®^ T7 Kit, Invitrogen, USA). The ability of T7 RP to incorporate different 5′-modified guanosine analogues was utilized in transcriptional priming to label and specifically conjugate the 5′-terminus of transcripts^4,5^. for incorporating a 5′-azide reactive group. In the IVT reaction, 5′-azido-5′-deoxy guanosine (5’-N_3_G) is added in fourfold excess over GTP to prime transcriptions ^1^. RNA formation was confirmed by gel electrophoresis (PAGE) (Figure S1). The transcript with the 5’-terminal azide (R-N_3_) is subsequently used in click reactions with a fluorophore or DNA strand containing a dibenzocyclooctyl (DBCO) group. The click conjugation is performed in 10 mM sodium acetate buffer pH 5.5. This conjugated strand was then annealed to the corresponding complementary strands heating at 90°C for 5 min, and then slowly cooling to room temperature at 5°C per 15 min.

A sample of duplex DNA for screening of DNA binding V_H_H was made by mixing the relevant DNA oligonucleotides in equimolar ratios, heating at 90°C for 5 min, and then slowly cooling to room temperature at 5°C per 15 min. This sample preparation was carried out with oligonucleotide concentrations of 5 μM, in phosphate buffered saline (PBS) of pH 7.3, in the presence of 100 mM KCl. Samples were then equilibrated at 4°C overnight. Samples were used within 7 days of annealing.

Selective oligonucleotides were phosphorylated at their 5’ end by incubating them with T4 polynucleotide kinase (PNK; New England Biolabs, USA). 2 nmoles of the oligonucleotide were mixed with 2 μL of 10× PNK buffer, 2 μL of PNK (10 U/μL), 4 μL of 1 mM ATP and the volume was made up to 20 μL. This reaction mix was incubated at 37°C for 1 h. Post-incubation, the enzyme was inactivated by incubating the mixture at 75°C for 15 min. The DNA was subsequently ethanol precipitated, re-suspended in Milli-Q water and quantified using UV absorbance. These were then used to prepare duplex DNA using the protocol mentioned above.

A sample of the neuronal targeting probe (nD^A647N^) was made by mixing the DNA oligonucleotides nD and nD’ in equimolar ratios, heating at 90°C for 5 min, and then slowly cooling to room temperature at 5°C per 15 min. This sample preparation was carried out with oligonucleotide concentrations of 5 μM, in 10 mM phosphate buffer of pH 7.4, in the presence of 100 mM KCl. Samples were then equilibrated at 4°C overnight.

### Preparation of helper phages

*Escherichia coli* strain TG1 grown in minimal media (M9 media) was inoculated 1:100 in 100 mL 2×TY media and grown till OD_600_ reached 0.2. 200 μL of this culture was then transferred to tubes and infected with serial dilutions of 4×10^11^ M13KO7 helper phage (with a minimum ratio of 1 bacterium per 20 helper phages, GE Healthcare, USA) (10-10, 10^−11^, 10^−12^, 10^−13^, 10^−14^). The tube was quickly transferred to a 37°C water bath and incubated for 30 min without shaking. 3 mL of H-Top agar was heated to 42°C and quickly poured to each tube. This was then poured on 2×TY plates and incubated overnight at 37°C. The plaques obtained from each plate were inoculated in 3 mL of 2×TY and grown till optical density at 600 nm (OD_600_) reaches 0.5. At this stage, the cultures were again infected with helper phage and incubated for 2 h at 37°C with shaking. This was diluted in 500 mL of 2×TY media and incubated at 37°C for 1 h, after which kanamycin was added to a final concentration of 50 μg/mL and the culture was incubated overnight at 37°C. The overnight culture was centrifuged at 10800×g for 15 min and the supernatant was collected into a glass conical flask. This flask was transferred to ice and 100 mL ice cold polyethylene glycol/NaCl solution (30% PEG 8000 with 2.5 M NaCl) was added with constant shaking. The resulting mixture was kept on ice for 45 min. The solution became turbid and was centrifuged at 10800×g for 30 min and the supernatant was discarded. The phage precipitate was re-suspended in 6 mL PBS containing 30% glycerol. This was filtered using a 0.45 μm membrane filter (Merck Millipore, USA), aliquoted and stored at −80°C.

### Preparation of library

A 250 μL aliquot of the glycerol stock library (4×10^10^ clones/mL) was inoculated in 250 mL of 2xTY containing 100 μg/mL of ampicillin (Sigma-Aldrich, USA) and 1% glucose. This was incubated at 37°C till the OD_600_ reached 0.5. 25 mL (containing 10^10^ clones) of this culture was infected with an excess of helper phage M13KO7, with a minimum ratio of 1 bacterium per 20 helper phages. Care was taken that no pipetting or shaking was done. This was incubated for 30 min at 37°C, without agitation, in a water bath. The infected bacteria were centrifuged for 20 min at 4200 rpm, the pellet was re- suspended in 500 mL of 2xTY with 100 μg/mL ampicillin and 50 μg/mL kanamycin without glucose, and incubated overnight at 30°C with shaking. The overnight culture was centrifuged at 10800×g at 4°C for 10 min. 100 mL of ice cold PEG-NaCl solution was added to the supernatant of the centrifuged culture, such that the supernatant becomes cloudy. This was incubated for 1 h on ice at 4°C. The mixture was centrifuged for 30 min at 10800×g at 4°C to pellet the phages. The PEG/NaCl solution was aspirated off carefully without disturbing the pellet. The pellet was re-suspended in 40 mL Milli-Q water, after which 8 mL of PEG/NaCl was added. The solution was swirled for efficient but gentle mixing and then allowed to stand on ice at 4°C for 20 min. The phages were centrifuged again at 10800×g for 30 min. The PEG/NaCl supernatant was carefully removed. The pellet containing a purified library of phages was re-suspended in 5 mL cold PBS and centrifuged again for 10 min at 13000 rpm at 4°C to pellet cell debris and bacteria. This is used as input for the phage display screen. The rest is stored as aliquots at −80°C.

### Preparation of DNA conjugated magnetic beads

150 μL streptavidin-coupled magnetic Dynabeads (Invitrogen, USA) were washed 3 times in PBS supplemented with 0.1% Tween-20 (Sigma-Aldrich, USA) (PBST). After each wash, beads were collected using a magnet. 15 μL of the B-RO3-CELL duplex (oligonucleotides Biotin-R-CELL+O3-CELL) was added to the washed magnetic beads and the volume of the reaction made up to 500 μL with PBST, such that the final DNA concentration used for the screening is 50 nM. This was incubated on a rotator for 1 h. After 1 h, the DNA coated beads are washed in PBST 3 times and then suspended in 150 μL of PBS. For each round of selection, 50 μL aliquots of this DNA conjugated magnetic bead mix were used. The rest was stored at −20°C for future use.

### Selection of dsDNA binders using phage display method

The phage display screen was carried out using a previously described protocol 10, with some variations. Non-specific binders (in particular anti-streptavidin V_H_Hs) were removed from the library by incubating V_H_Hs displayed on the surface of phages with streptavidin-coated magnetic beads alone. 50 μL of streptavidin coated magnetic beads were mixed with 100 μL of the library (this input contains 1.1×10^12^ phages) and incubated for 30 min on a roller and another 30 min standing. Volume of this mix was made up to 1.5 mL with PBST supplemented with 0.6% non-fat milk (PBSTM). After 1 h, the beads containing streptavidin binders were pulled down using a magnet, while the supernatant containing the remainder of the library was added to 50 μL of the DNA coated magnetic beads. This mix was incubated on a roller for 30 min and then left standing for 1.5 h. After 2 h, the beads were pulled down using a magnet, the PBSTM was removed, 1 mL of fresh PBSTM was added and the beads were fully re-suspended. This mix was added to a 15 mL polypropylene tube, which was pre-blocked with PBSTM (2% non-fat milk), and the volume made up to 10 mL using PBST. The beads were collected using a magnet for 5 min, the PBST was removed and fresh PBST was added. This was repeated 20 times, with a change of polypropylene tube every 5 washes. After the last wash, 1 mL of 100 mM triethylamine (TEA) was added to re-suspend and dissociate the phages from the beads; this was transferred to a tube and rocked for 7 min, after which the beads were collected using a magnet. 500 μL of the supernatant was carefully added to a 500 μL of 1 M Tris-Cl (pH 7.4) to neutralize it; the remaining 500 μL was put back on the roller again for 7 min and the above steps were repeated. Finally, 200 μL of 1 M Tris-Cl was added to the polypropylene tube and kept separately.

### Rescue and enrichment of possible DNA binders

The rescue and amplification of selected phage was done by infecting *E. coli* TG-1 bacteria with eluted phage. 750 μL of the recovered phages were added to 9.25 mL of a TG1 culture (grown to an OD_600_ of 0.5 in 2×TY media with 1% glucose) and incubated at 37°C for 30 min in a water bath without agitation. Simultaneously, 4 mL of the same TG1 culture was added to the last polypropylene tube and the infection protocol was repeated. Both the cultures were pooled to get 14 mL of infected TG1 culture. Of this, 3 dilutions of 1 mL each were made, 10^−1^, 10^−2^ and 10^−3^, and two volumes of each, 10 μL and 100 μL, were spread on 2×TY plates containing 1% glucose and 100 μg/mL ampicillin. These plates were used for calculation of phage output after the first round of selection. The remaining culture was spun down at 3300 rpm for 10 min at RT. The pellet was re-suspended in 1.5 mL 2×TY, which was equally divided and plated on 3 large 2×TY plates containing 1% glucose and 100 μg/mL ampicillin. All plates were grown overnight at 37°C.

Colonies obtained on the large petri dishes were scraped off using 6 mL of 2×TY supplemented with 30% glycerol. This scraped culture constitutes the input for Round 2 of screening. An aliquot of this culture was added to 100 mL of 2×TY with 1% glucose and 100 μg/mL ampicillin, such that OD_600_ of the inoculated culture was 0.05. This was incubated at 37°C till OD_600_ reaches 0.5. 10 mL was aliquoted into a fresh tube, helper phages were added (bacteria: helper phage = 1:20; at OD_600_ = 0.5, a 10 mL culture of *E.coli* contains 4×109 bacteria) and the culture was incubated at 37°C in a water bath for 30 min without agitation. The culture was centrifuged at 3300 rpm for 10 min at RT, the pellet was re-suspended in 50 mL 2×TY with 100 μg/mL ampicillin and 50 μg/mL kanamycin (without glucose) and incubated overnight at 30°C.

40 mL of the overnight culture was centrifuged at 10800×g at 4°C for 10 min. 8 mL of ice cold PEG-NaCl solution was added to the supernatant of the centrifuged culture, such that the supernatant becomes cloudy. This was incubated for 1 h on ice at 4°C. The mixture was centrifuged for 10 min at 10800×g at 4°C to pellet the phages. The PEG solution was aspirated off carefully without disturbing the pellet. The pellet was re-suspended in 2 mL cold PBS without introducing air bubbles. The phages were centrifuged again at 10800×g for 10 min. The supernatant was carefully removed. The pellet containing cell debris and bacteria was discarded, while the supernatant was used as input for Round 2 of selection. 500 μL dilutions of 10-9, 10-10 and 10-11 were also made with the recovered phages, introduced into TG1 bacteria (as described above) and spread on 2×TY plates containing 1% glucose and 100 μg/mL ampicillin to calculate input. This was done for 3 rounds of selection. Negative selection against streptavidin was done at each stage of selection to completely eliminate anti-streptavidin V_H_Hs.

After 3 rounds of screening, 80 clones each from Round 2 and 3 were selected for further characterization. Individual colonies were transferred to one well of a 2 mL deep 96-well plate (Greiner Bio-one, Germany) with 600 μL 2×TY containing 100 μg/mL ampicillin and 1% glucose, and grown overnight with shaking at 37°C. Glycerol was then added to a final concentration of 30% to make a master plate which was stored at −80°C.

### ELISA using phages

ELISA was performed using phages secreted in the media, each of which carries the V_H_H fused to a coat protein. 600 μL of 2×TY with 100 μg/mL ampicillin and 1% glucose were added to each well of 2 mL deep 96 well plates, each of which was inoculated with 6 μL of the master stock. This was incubated for 2.5 h at 37°C with shaking, such that OD_600_ reaches 0.5. Helper phages were added to each well and the plates were incubated at 37°C without agitation. The plates were then centrifuged at 2500 rpm for 5 min. The supernatant was carefully aspirated off and the pellet re- suspended in 600 μL 2×TY containing 100 μg/mL ampicillin and 50 μg/mL kanamycin. The plates were incubated overnight at 30°C with agitation. The phages were recovered by centrifuging the cultures at 2800 rpm for 10 min at RT and pipetting the supernatant in fresh 96 well plates.

In order to perform the ELISA, 96 well ELISA plates (Nunc Maxisorp, Thermo Fisher Scientific, USA) were coated with 50 μL of 20 μg/mL avidin and incubated at RT for 2 h. The plates were flicked and washed once with PBS to remove excess avidin. 50 μL of biotinylated DNA at 50 nM concentration was immobilized by adding it to the wells and incubating at 4°C overnight. Excess DNA was removed by flicking the plates. The plates were blocked using 200 μL of and kept at RT for 1 h, after which it was removed by flicking. In a separate plate, 90 μL of phages were mixed with 20 μL of PBSTM, incubated at RT for 20 min and then 100 μL of this mix was added to the ELISA plate. The phages and DNA were allowed to bind for 2 h, after which the plates were flicked and washed 3 times each with PBST and PBS. 50 μL of anti-M13 antibody conjugated to horseradish peroxidise (HRP; GE Healthcare Life Sciences, USA) was added at a 1:5000 dilution and incubated for 40 min. The plates were flicked and washed 3 times with PBST and PBS. 100 μL of tetramethyl benzidine (TMB)/H_2_O_2_ (BD Biosciences, USA) was added to each well, and the reaction was stopped by addition of 100 μL of 2 N H_2_SO_4_. Binding was quantified by measuring absorbance at 450 nm using a Spectramax multi-mode plate reader (Molecular Devices, USA).

### ELISA using secreted V_H_Hs

ELISA was performed using V_H_Hs secreted in the media. 600 μL of 2×TY with 100 μg/mL ampicillin and 1% glucose was added to each well of 2 mL deep 96 well plates, each of which was inoculated with 6 μL of the master stock. This was incubated for 2 h at 37°C with shaking, after which 1 mM isopropyl thiogalactoside (IPTG) was added. The plates were then shifted to 30°C and incubated overnight. The cultures were centrifuged at 2800 rpm for 10 min at RT and the supernatant, which contains secreted V_H_Hs, was carefully transferred to a fresh 96 well plate. After this, standard ELISA, as outlined in Section IV.B.8.1 was carried out, using mouse anti-Myc antibody (Sigma-Aldrich, USA) at a dilution of 1:1500 and goat anti-mouse antibody conjugated to HRP (Life Technologies, USA) at a dilution of 1:1000.

### Characterization of sequence specificity

Approximately 80 colonies each from Round 2 and Round 3 of selection were screened by ELISA against the dsDNA of choice (Biotin-R-CELL+O3-CELL). This screening was done using both phage and protein ELISA, as described above. In order to check the sequence specificity of the positive clones, V_H_Hs were subjected to another round of ELISA assay against various DNA epitopes. The epitopes used were ssDNA (Biotin-R-CELL, Biotin-O3-CELL), dsDNA (Biotin-R-CELL+O3-CELL), divergent dsDNA (Biotin-DS1-CELL+DS2-CELL) and various parts of the dsDNA (Biotin-R-CELL+R1-CELL, Biotin-R-CELL+RM-CELL, Biotin-R-CELL+R2-CELL, Biotin-R-CELL+R3-CELL, Biotin-R-CELL+R4-CELL, Biotin-R-CELL+R5-CELL). After immobilization of these targets, ELISA assay was carried out as mentioned above.

### Sequencing of specific clones

About 42 clones showing binding to dsDNA were chosen for sequencing. Individual clones were grown overnight and subjected to a plasmid DNA isolation using Nucleospin Plasmid Miniprep Kit (Macherey-Nagel GmbH, Germany). Sequencing was performed using a standard dideoxy sequencing method.

### Expression and purification of V_H_Hs

V_H_H expression was performed in M9 minimal media (1× M9 salts, 2 mM MgSO_4_, 1% glycerol, 0.1% casamino acids and 0.000005% thiamine). Selected V_H_H was inoculated in 100 mL 2×TY supplemented with 100 μg/mL ampicillin and 1% glucose and grown overnight at 37°C. 5 ml of this culture was inoculated in 500 mL of M9 minimal media with 100 μg/mL ampicillin and grown at 37°C for 2 h. IPTG was added to a final concentration of 1 mM, after which the culture was shaken at 30°C for 16 h. Bacteria were centrifuged at 10000 rpm for 10 min. The supernatant containing secreted V_H_Hs was filtered using a 0.22 μm membrane filter to remove the remaining debris. The filtered media was then incubated with 2 mL of Talon Cobalt affinity resin (Clontech Laboratories Inc., USA) for 1 h at 4°C. Post-binding, the media with the resin was put into a funnel adapted onto a column, which was pre-washed with 20 mL of PBS, at 4°C. The flow through was collected and again passed through the column in order to collect all the metal beads with bound protein. This was done 3 times. Once all the beads were collected, they were washed again by passing 100 mL of PBS through the column. Non- specific binding to beads was abrogated using 1 mL 5 mM imidazole. The bound V_H_H was then eluted using a gradient of imidazole concentrations, ranging from 50 mM to 250 mM. The first 9 fractions were run on a 12% SDS-PAGE to check the expression and elution of the protein.

Each eluted V_H_H was further purified by removing imidazole using a 10 kDa Amicon ultra centrifugal filter (Merck Millipore, USA). Each fraction was added to the centrifugal filter and centrifuged for 10 min at 4°C for 14000 rpm. Volumes were then made up to 400 μL using PBS after each spin. This was repeated 10 times. Concentration of each fraction was measured using Bradford assay. Purified V_H_H were stored at 4°C for short term and at −20°C for long term.

### pH dependent ELISA

pH dependent ELISA to check pH sensitivity of the selected V_H_H was performed using purified V_H_H. The V_H_H solution was divided into two pools and each pool was incubated with PBSTM of the desired pH for 20 min. This was then added to ELISA plates containing the immobilized DNA of choice. Standard ELISA, as described above, was then employed to assess binding.

### Determination of V_H_H binding affinities

Affinity of selected V_H_Hs for their dsDNA epitopes was assessed using 3 formats, as described below. All data was analyzed using OriginPro 8.5 (OriginLab, USA).

#### i) Using serial dilutions of immobilized DNA

Avidin coated 96-well plates were incubated with serial dilutions of the dsDNA antigen (10 nM - 5 μM), as described in Section IV.B.8. 250 nM of the protein was added and allowed to bind for 2 h. ELISA was then carried out as described above.

#### ii) Using serial dilutions of competitive DNA

500 nM of the biotinylated dsDNA antigen was mixed with serial dilutions of competitor non-biotinylated dsDNA (1 nM - 25 μM), to which 100 nM of protein was added. This mixture was allowed to incubate for 2 h at RT, after which it was added to avidin coated 96-well plates and further incubated for 2 h. ELISA was then performed.

#### iii) Using serial dilutions of purified V_H_H

Avidin coated 96-well plates were incubated and bound with 500 nM of the dsDNA antigen as described in Section IV.B.8. Serial dilutions of the protein (1 nM - 5 μM) were added and allowed to bind for 2 h. ELISA was then carried performed.

### Electrophoretic mobility shift assay

The required dsDNA constructs were annealed, as described above, at concentrations of 5 μM. A binding reaction was set up consisting of 1 μM DNA and 1 μM protein in PBS. Glycerol was added to a final concentration of 10% in order to minimize DNA-protein complex dissociation. DNA-protein complexes were allowed to form at RT for 2 h, after which 50 pmoles of DNA were loaded on an 8% native PAGE. The gel was run at 4°C using Tris-acetate-EDTA (TAE) buffer at 100 V. The DNA-protein complexes were visualized using ethidium bromide staining.

### Plasmid vectors and construction of 9E fusions

PHA-1 plasmid (gift from Krastosis lab, University of Chicago) and pBluescript SK(-) (Agilent Technologies, USA) were used during construction of transgenic strains. PCR fragments containing promoters of choice, *snb-1* or *odr-2* and 9E were generated using standard cloning protocols. *psnb-1:odr-2* and *snb-1::9E* geneBlock fragments were obtained from IDT. *pglr-4, psnb-1, 9E* was cloned out of plasmids generated in house (*pglr-4-* Krastosis lab; *psnb-1* and 9E*-* Krishnan lab). *pacr-2 was* cloned out of genomic DNA isolated from wild type worms. All the PCR fragments had a *unc-54*-3’UTR sequence on their 3’ end for better expression in worms^6^.

### *C. elegans* methods and strains

Wild type strain used was the *Caenorhabditis elegans* isolate from Bristol (strain N2)^7^. Mutant strains used are VC1119 (*dyf-2&ZK520.2(gk505) III*), VC1521 (*dyf-2(gk678) III*), HC196 (*sid-1(qt9) V*). Other transgenics used for colocalization studies are as follows

1. *jsIs682* [p*rab-3::gfp::rab-3*]^8,9^ expresses GFP::RAB-3 in all neurons.
2. *otIs355* [*rab3p(prom1)::2xNLS::TagRFP*] which expresses RFP in the nucleus of all neurons^10^
3. *otIs45* [*unc-119::GFP*] which expresses GFP in the cytosol of all neurons^11^
4. *pwIs50* [*lmp-1::GFP + Cbr-unc-119(+)*] which expresses LMP-1::GFP a lysosomal marker^12^.
5. *hjIs9* [*ges-1p::glo-1::GFP + unc-119(+)*] in which GFP is targeted to lysosome related organelles (LROs) in intestinal cells^13^

All strains were procured from the *Caenorhabditis* Genetics Centre (CGC; University of Minnesota, USA). Standard methods were followed for the maintenance of *C. elegans*.

Transgenic strains were created by co-injecting PCR fragment of the gene of interest and PHA-1 plasmid (20 ng/μl) as a selectable marker in *pha-1* mutants^14^. The PCR fragments used in this study were *psnb-1:snb-1::9E psnb-1:odr-2::9E; pglr-4:snb-1::9E* and *pacr-2:snb-1::9E.* Total concentration of injected DNA was 100 ng/μl, which was achieved using empty pBluescript SK(-) vector. Transgenic lines obtained were also maintained using standard protocols. *pha-1* mutants are temperature sensitive embryonic lethal mutants which can grow at 15°C but not 25 °C^15^. Each line was characterized based on the survival at 25 °C.

### Intestinal epithelia and neuron labelling

Intestinal labeling was done by soaking worms in a 10 μL, 1 μM solution of intestine targeting probe at pH 5.0 for 2 hours. Worms were then washed in PBS and placed on OP50 plates for another hour before imaging. Worms were mounted on 2.0% agarose pads and anesthetized using 40 mM sodium azide in M9 buffer.

Neuronal labelling was done by injecting 500 nM of nD^A647N^. Injections were performed, as previously described^16^, in the dorsal side in the pseudocoelom, just opposite the vulva, of one-day old adult wild type hermaphrodites. Injected worms were mounted on 2.0% agarose pads and anesthetized either using 40 mM sodium azide or 1mM levimasole in M9 buffer. Neuronal labeling was examined after 30-60 minutes of incubation at 22°C. Imaging of labelled neurons was carried out in at least 15 worms.

### Co-localization experiments

For gut colocalization experiments ~10 one-day adult transgenic worms were incubated in a solution of 1 μM R^100^D^38^ for two hours. Worms were then washed and incubated on OP50 plates for 1 hour for the clearing of the intestine lumen of excess sensor.

For neuronal co-localization experiments *psnb-1:snb-1::9E* were crossed with various transgenic lines containing fluorescently labelled neuronal markers. Briefly, L4 for hermaphrodite worms of various fluorescently labelled transgenic lines were crossed with N2 males and the male progeny containing a fluorescent label were selected for the next step. Fluorescently labeled male worms were crossed with *psnb-1:snb-1::9E* containing L4 hermaphrodites. 10-12 hermaphrodite progeny from this step were singled out at the L4 developmental stage based on the presence of a fluorescent marker. These worms were allowed to self-fertilize and 5 of their progenies were used to perform single worm PCRs to identify which plates contain worms with both 9E and the fluorescent neuronal markers. Single worm PCRs were performed using standard methods^17^. Briefly, 5 worms from each plate were placed into a tube containing 5μL of worm lysis buffer (WLB: 50 mM KCl, 10 mM Tris pH 8.3, 2.5 mM MgCl_2_, 0.8% Tween-20, 0.01% Gelatin) and 100 μg/ml proteinase K. Worms are frozen at −80°C for at least one hour. The tube is immediately treated at 60°C for 60min after which proteinase K is inactivated by heating to 95°C for 15min. The PCR mix is then added directly to the digested heat inactivated sample. The PCR product was then confirmed by gel electrophoresis for 9E. Actin was used as a control. Plates which showed the presence of *psnb-1:snb-1::9E* were taken forward for injecting the DNA device similar to protocols mentioned above.

Worms were mounted on 2.0% agarose pads and anesthetized using either 40 mM sodium azide or 1 mM levimasole in M9 buffer and imaged on Leica TCS SP5 II STED laser scanning confocal microscope (Leica Microsystems, Inc., Buffalo Grove, IL, USA) in confocal mode.

### Microscopy and image analysis

Confocal images were captured with a Leica TCS SP5 II STED laser scanning confocal microscope (Leica Microsystems, Inc., Buffalo Grove, IL, USA) equipped with 63X, 1.4 NA, oil immersion objective. Alexa 488 was excited using an Argon ion laser for 488 nm excitation, Alexa 647 using He-Ne laser for 633 excitation. Images on the same day were acquired under the same acquisition settings. All the images were background subtracted prior to any image analysis, which was carried out using ImageJ ver.1.49d (NIH, USA).

Image analysis for quantification of uptake during intestinal labeling was done using custom MATLAB code. For each worm the most focused plane was manually selected in the Alexa 647 channel. To determine the location of the endosome first a low threshold was used to select the entire field. Only the area within the cell was subsequently considered for vesicle selection. Regions of interest corresponding to individual vesicle were selected in the Alexa 647 channel by adaptive thresholding using Sauvola’s method ^18^. The initial selection was further refined by watershed segmentation and size filtering. After segmentation regions of interest were inspected in each image and selection errors were corrected manually. Using this we measured the mean fluorescence intensity for ~500 vesicles from ~10 animals and the background intensity corresponding to that field was subtracted.

**Supplementary Fig 1:**
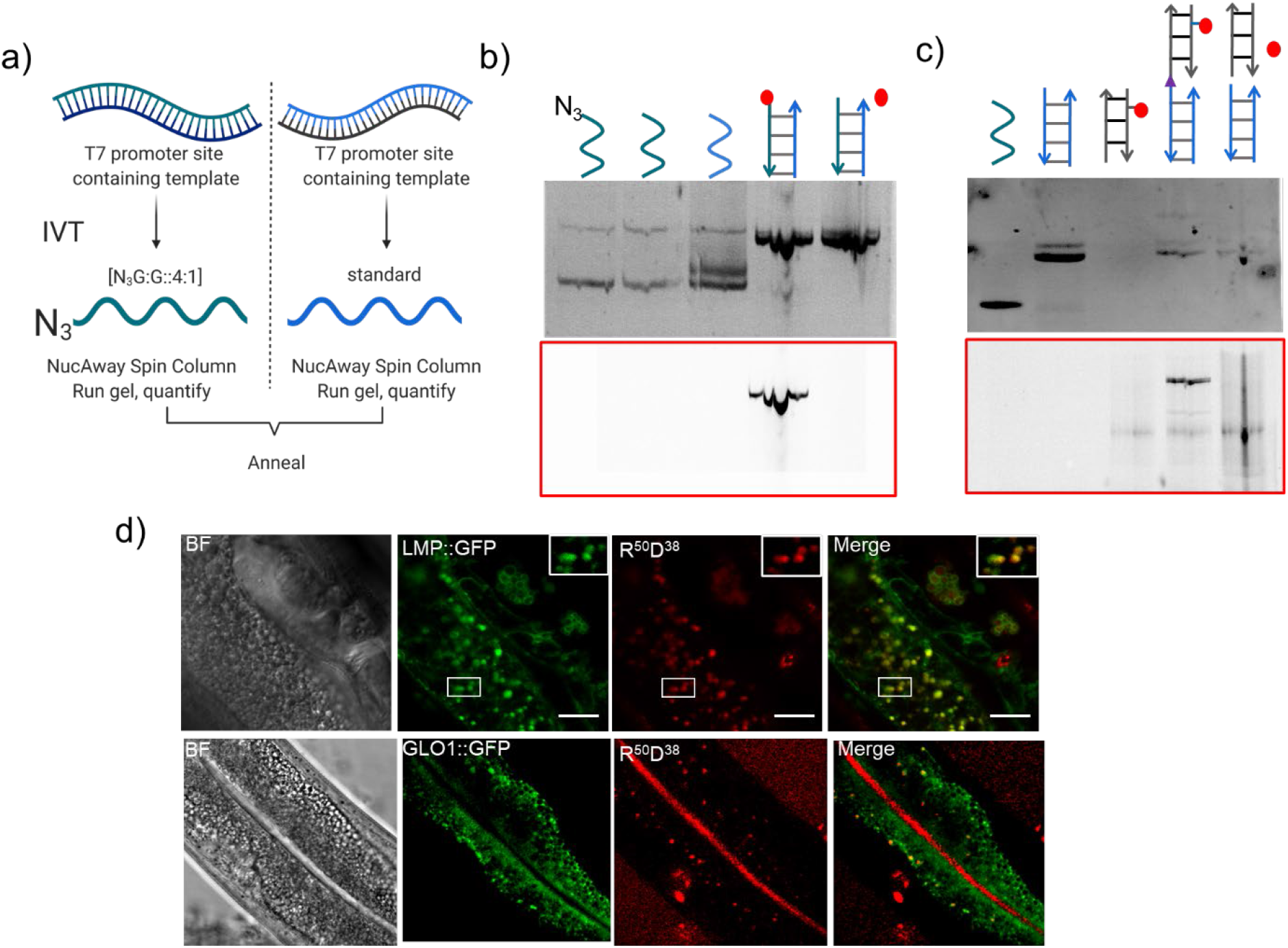
Uptake of nucleic acid probes in the intestinal epithelial cells. a) Schematic showing protocol of synthesis of azide labelled RNA for R^50^ and R^100^ strands. b) The conjugation of Alexa 647N was confirmed by gel electrophoresis for R^50^. The 8% Native PAGE gel was imaged in the EtBr channel and Alexa 647N channel (red box). c) The conjugation of R^100^ (lane 2) with D^38^ (lane 3) to form R^100^D^38^ (lane 4) was confirmed by gel electrophoresis. The 15% Native PAGE gel was imaged in the EtBr channel. d) Representative images of colocalization between LRO markers (LMP1::GFP and GLO-1::GFP) and R^100^D^38^ fluorophore labeled nucleic acid probes in intestinal epithelial cells.

To test whether our R^n^D^n^ devices localized in LROs we performed colocalization experiments in transgenic worms expressing fluorescently labelled lysosomal markers LMP1::GFP or GLO-1::GFP that label lysosomes and LROs respectively. The transgenics were incubated in a solution of 1 μM R^100^D^38^ for 2 h. Worms were then washed and incubated on OP50 plates for 1 hour for clearing their intestine lumens of excess R^100^D^38^. Our experiments revealed that R^100^D^38^ and LMP1::GFP showed high colocalization, indicating that the nanodevices labeled LROs. Similarly R^100^D^38^ and GLO-1::GFP also colocalized. However, there was lower uptake into the IEC’s and poorer clearance of the gut lumen in GLO-1::GFP transgenic worms.

---

**Supplementary Figure 2:**
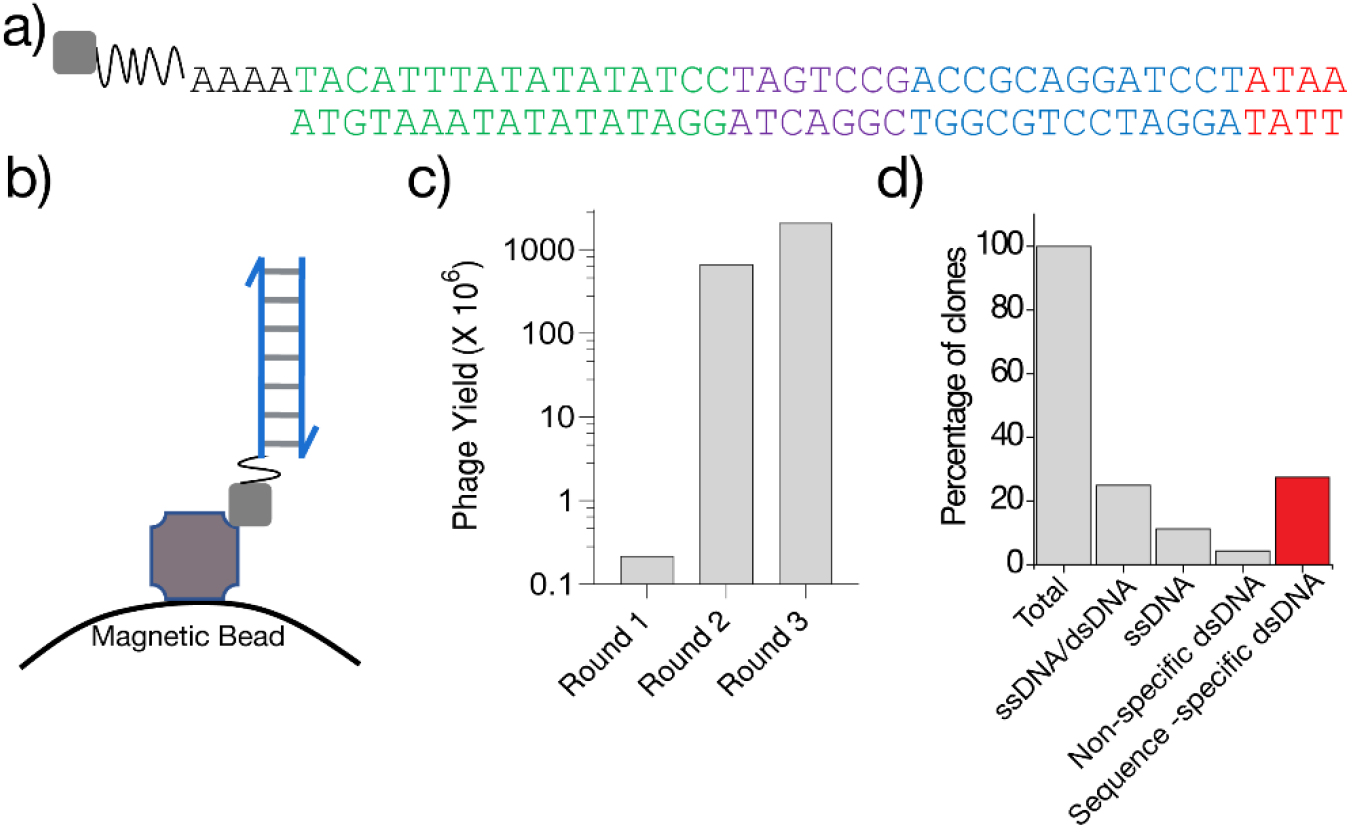
Analysis of recombinant antibody binders of a dsDNA epitope. a) Sequence of the biotin-labeled dsDNA epitope used for the phage display screen. b) The epitope was immobilized via biotin (grey square) on streptavidin conjugated magnetic beads. c) Yield of the phages after each round of selection, calculated as described in Materials and Methods. d) Percentage of V_H_H clones obtained that bound the indicated epitope and their sequence specificity.

**Supplementary Figure 3:**
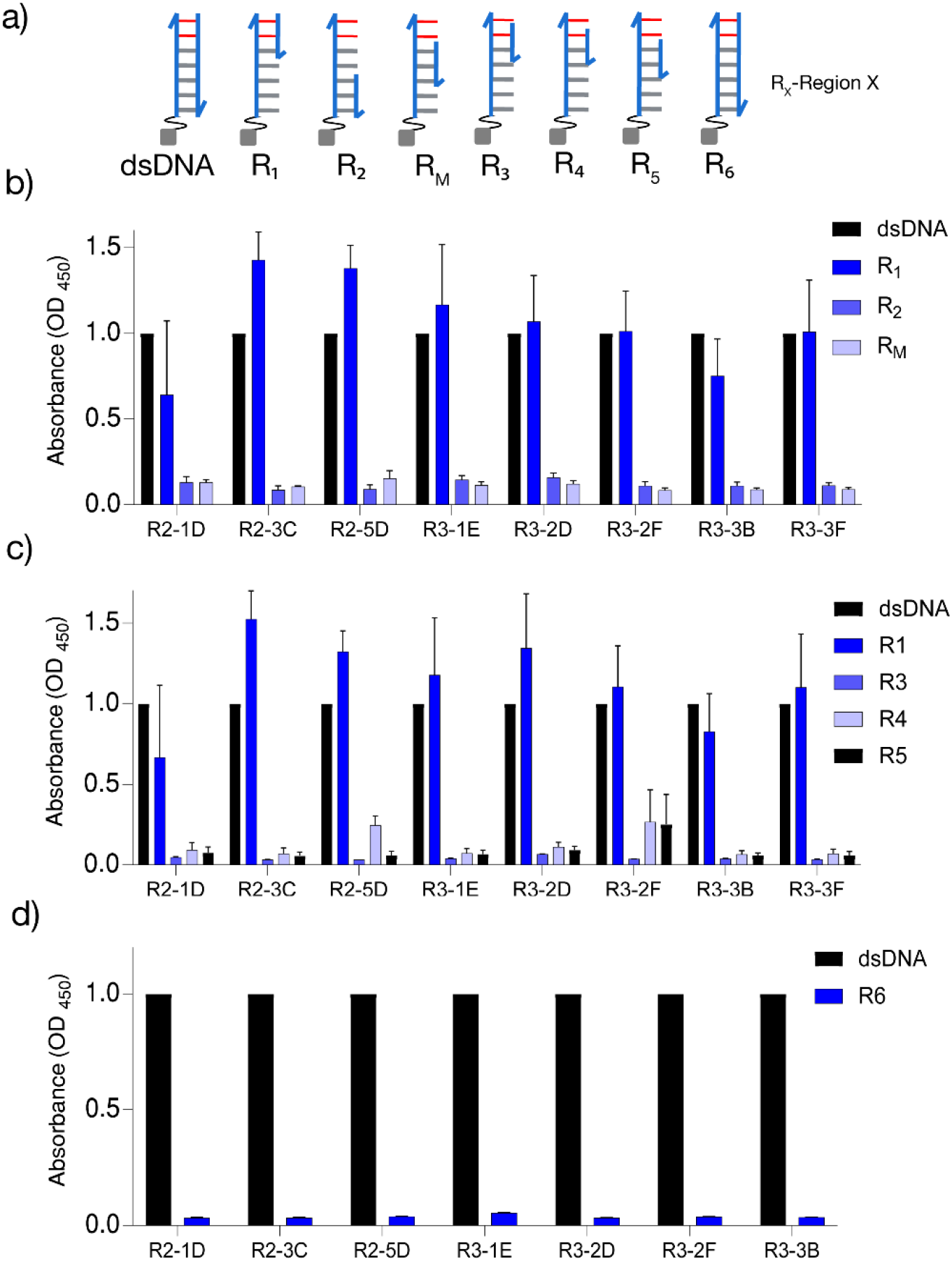
Characterization of dsDNA-binding V_H_H antibodies. a) Schematic of the dsDNA constructs used to find the minimal dsDNA-binding motif. All sequence details are provided in Supplementary Table 1. Epitopes were immobilized on streptavidin conjugated magnetic beads via biotin (grey square). Relative binding of a few representative dsDNA-binding V_H_H antibodies to b) duplexes R_1_, R_2_ and R_M_, c) duplexes R_3_, R_4_ and R_5_, d) duplex R_6_. All ELISA experiments were performed in triplicate and errors are presented as mean ± s.e.m.

**Supplementary Figure 4:**
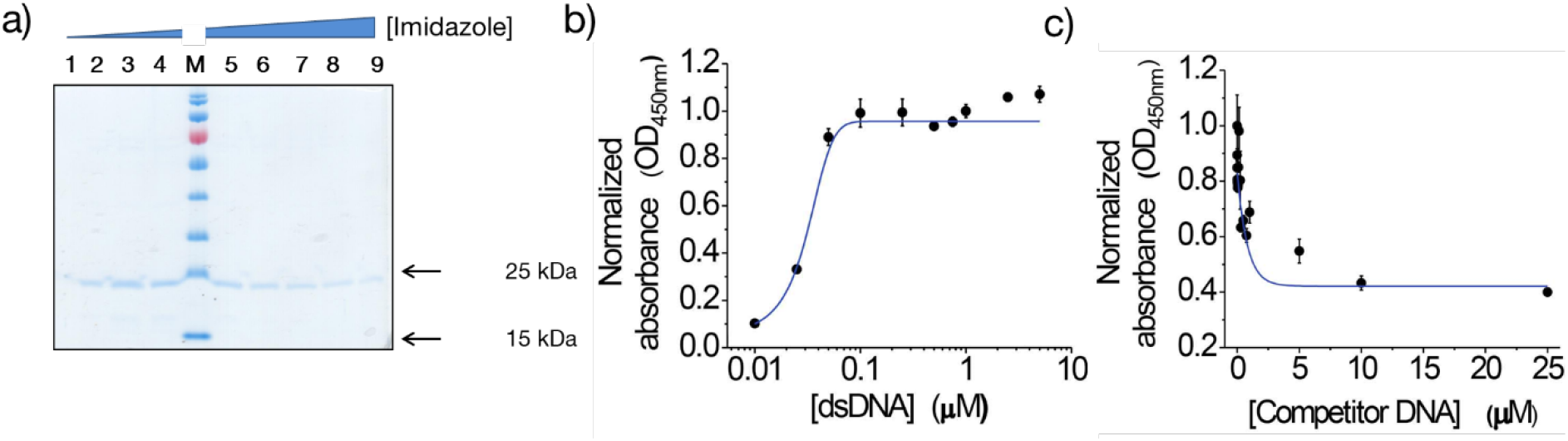
a) Purification of the dsDNA-binding 9E. Clone 9E was expressed and purified as described in Materials and Methods, and the fractions obtained after elution using increasing concentrations of imidazole were resolved on a 12.5% SDS-polyacrylamide gel. Fractions 3-9 were found to be enriched in the eluted protein. Semi-quantitative determination of affinity of 9E for the 41-mer dsDNA using (b) increasing concentrations of immobilized dsDNA and fixed amount of protein (250 nM), and (c) immobilized dsDNA (500 nM) in the presence of increasing amounts of competitor dsDNA. All experiments were performed in triplicate and the data is represented as mean ± s.e.m.

**Supplementary Figure 5:**
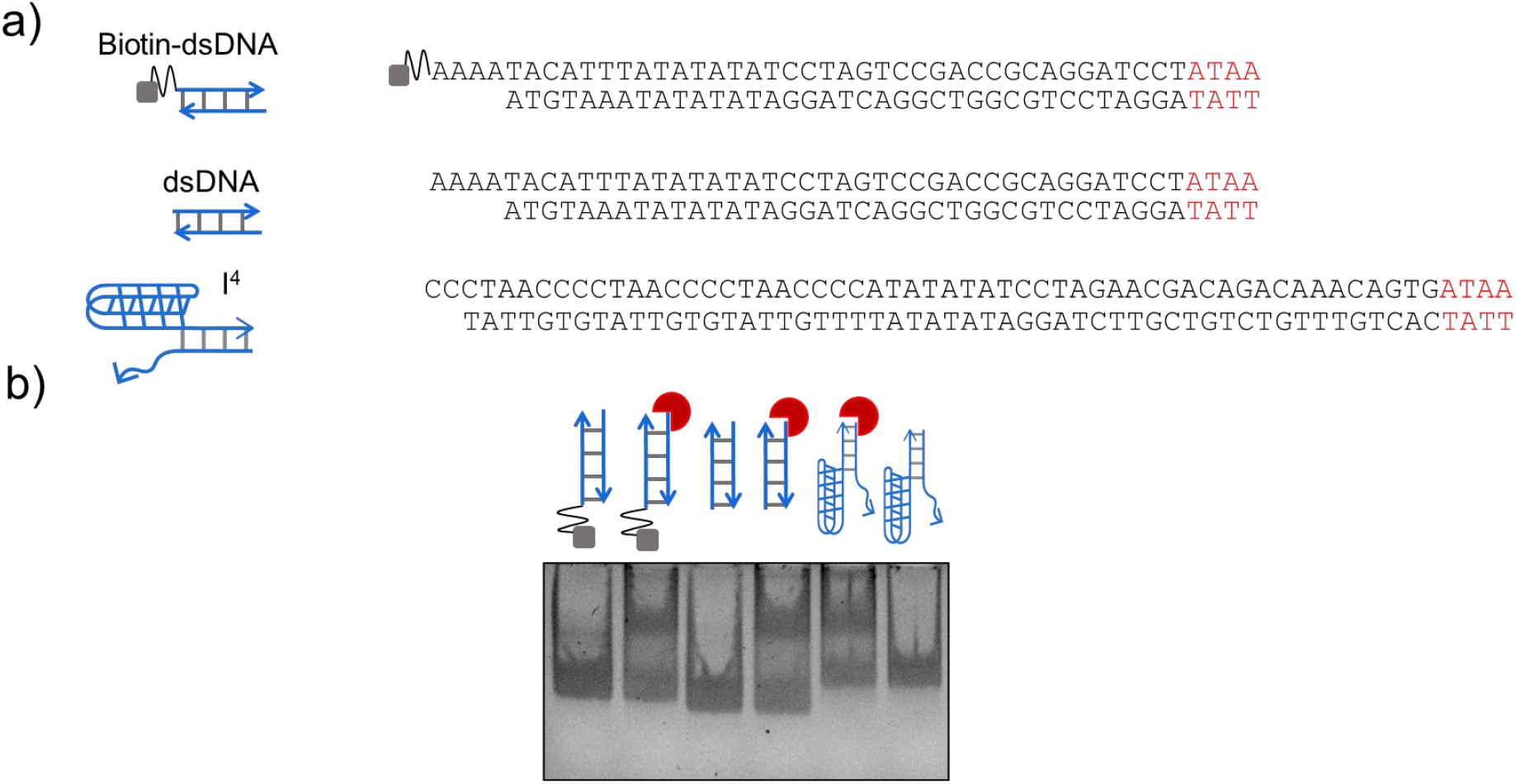
Electrophoretic mobility shift assay (EMSA) to demonstrate binding of 9E to the 4 nt minimal binding motif. (**a**) Sequences, and the corresponding schematics, of DNA constructs engineered with the 4 nt epitope (red) to demonstrate binding of 9E. **(b).**EMSA of 9E (red crescent) to biotinylated dsDNA (lanes 1, 2), dsDNA (lanes 3, 4) and a DNA duplex (I^4^; lanes 5, 6) where the 4 nt epitope was placed at one end of the duplex (region highlighted in red), while the other end comprised an I-motif forming C-rich segment. Binding assays were set up as described in Materials and Methods and run on an 8% non-denaturing polyacrylamide-TAE gel.

**Supplementary Figure 6:**
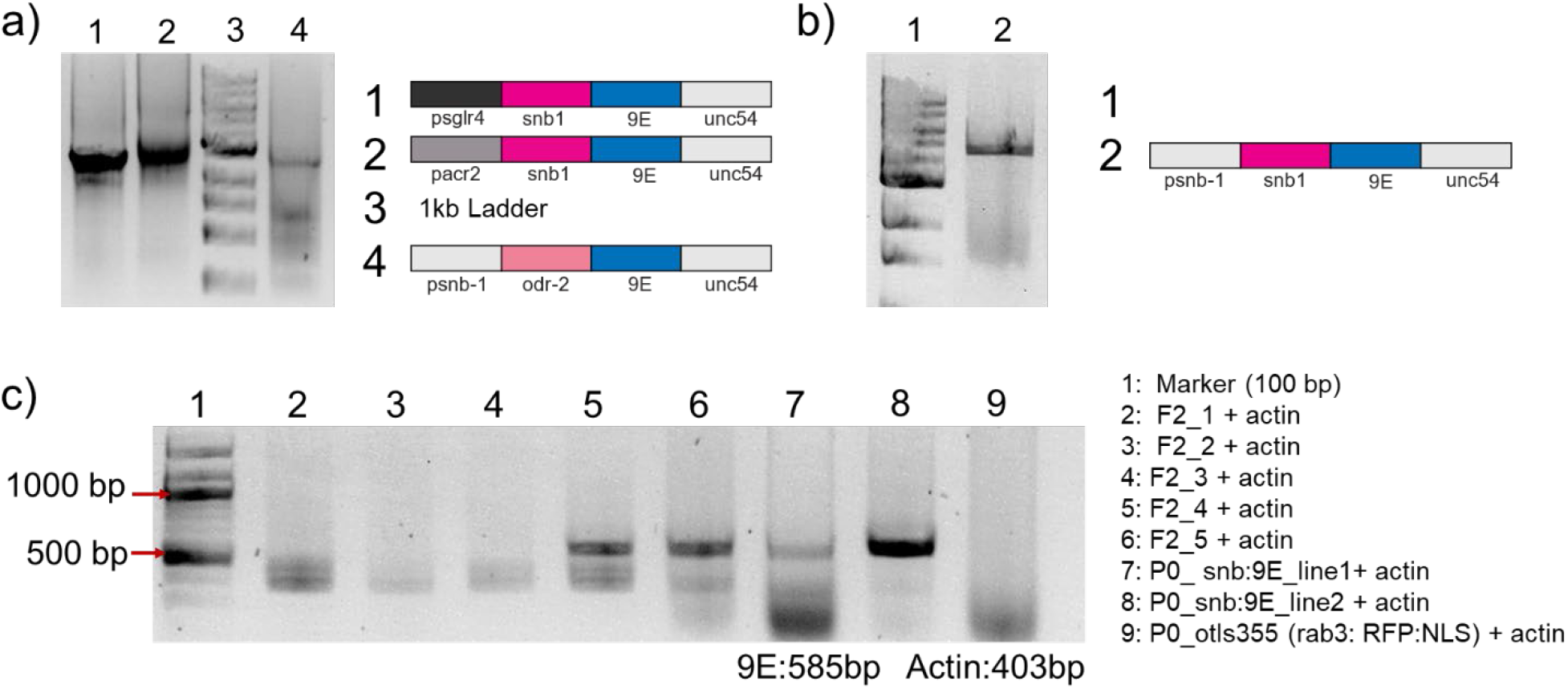
a) and b) PCR fragments used for making various transgenic strains to target DNA probes to neurons. PCR product was confirmed by gel electrophoresis with 1% Agarose gel and c) single worm PCR showing selection of worms containing p*snb-1:snb-1:: 9E* and 2XNLS::RFP

**Supplementary Figure 7:**
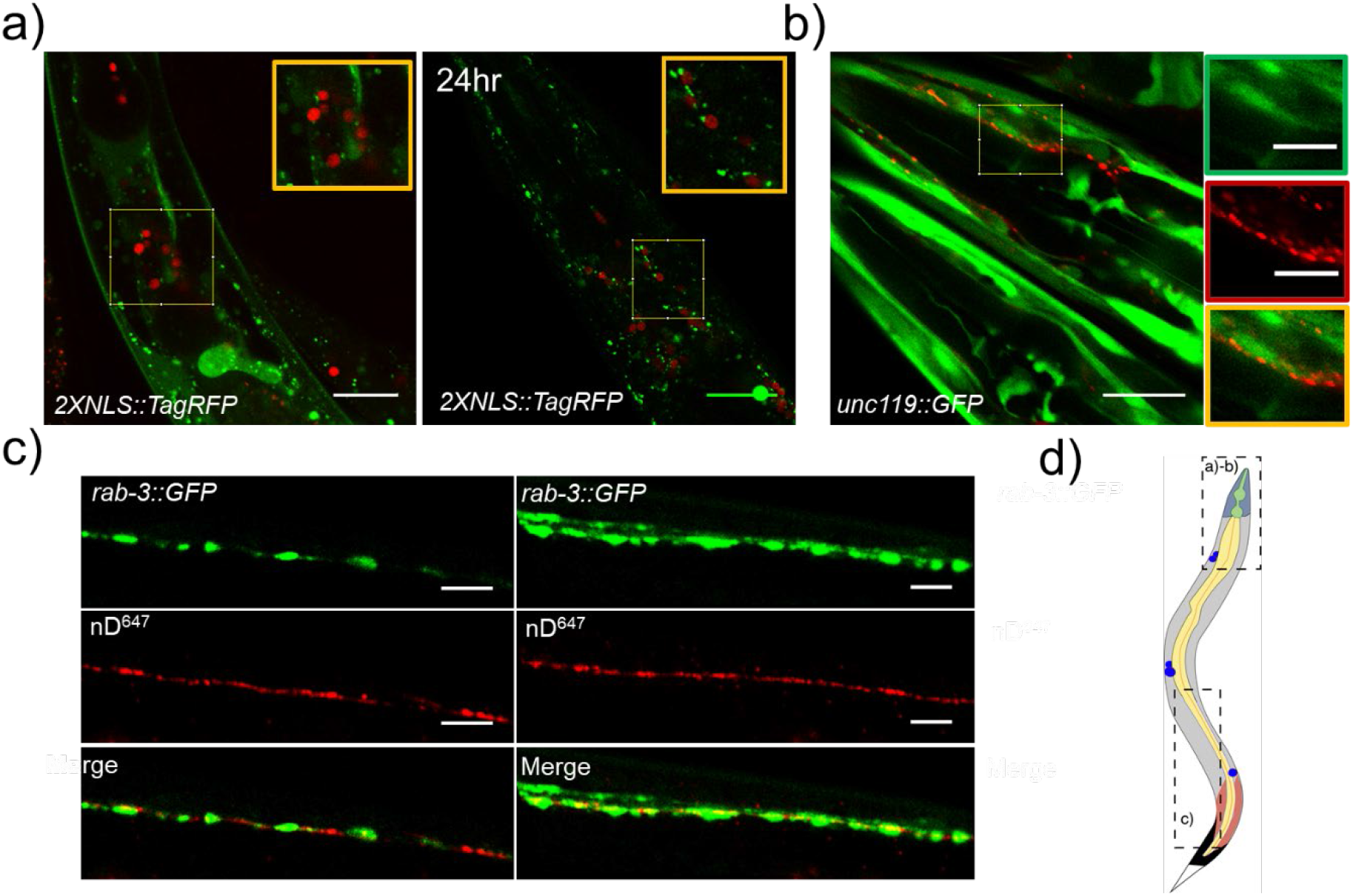
Schematic showing selection of worms containing psnb-1:snb-1:: 9E and 2XNLS::RFP: Representative merged images of transgenic worms containing both a) *rab-3p(prom1)::2xNLS::TagRFP* or b) *unc-119::GFP* or c) *prab-3::gfp::rab-3* and *psnb-1:snb-1::9E* injected with nDNA. d) Schematic showing regions of the worm represented in a-c. Scale bar a) 20 μm; b) 20 μm, inset-5μm and c) 5μm.

## Bibliography

Altun, Z. F., & Hall, D. H. (2009). Alimentary system, intestine. . WormAltas. Retrieved from http://doi:10.3908/wormatlas.1.4

Altun-Gultekin, Z., Andachi, Y., Tsalik, E. L., Pilgrim, D., Kohara, Y., & Hobert, O. (2001). A regulatory cascade of three homeobox genes, ceh-10, ttx-3 and ceh-23, controls cell fate specification of a defined interneuron class in C. elegans. Development, 128(11), 1951–1969.

Antibodies, M., Phage, B. Y., & Technology, D. (1994). MAKING ANTIBODIES BY PHAGE.

Bhatia, D., Arumugam, S., Nasilowski, M., Joshi, H., Wunder, C., Chambon, V., … Krishnan, Y. (2016). Quantum dot-loaded monofunctionalized DNA icosahedra for single-particle tracking of endocytic pathways. Nature Nanotechnology, 11(12), 1112–1119. doi:10.1038/nnano.2016.150

Bhatia, D., Surana, S., Chakraborty, S., Koushika, S. P., & Krishnan, Y. (2011). A synthetic icosahedral DNA-based host-cargo complex for functional in vivo imaging. Nature Communications, 2, 339. doi:10.1038/ncomms1337

Brockie, P. J., Madsen, D. M., Zheng, Y., Mellem, J., & Maricq, A. V. (2001). Differential expression of glutamate receptor subunits in the nervous system of Caenorhabditis elegans and their regulation by the homeodomain protein UNC-42. The Journal of Neuroscience, 21(5), 1510–1522.

Bujold, K. E., Lacroix, A., & Sleiman, H. F. (2018). DNA Nanostructures at the Interface with Biology. Chem, 4(3), 495–521. doi:10.1016/j.chempr.2018.02.005

Chakraborty, K., Leung, K., & Krishnan, Y. (2017). High lumenal chloride in the lysosome is critical for lysosome function. eLife, 6, e28862. doi:10.7554/eLife.28862

Chakraborty, K., Veetil, A. T., Jaffrey, S. R., & Krishnan, Y. (2016). Nucleic Acid-Based Nanodevices in Biological Imaging. Annual Review of Biochemistry, 85, 349–373. doi:10.1146/annurev-biochem-060815-014244

Chou, J. H., Bargmann, C. I., & Sengupta, P. (2001). The Caenorhabditis elegans odr-2 gene encodes a novel Ly-6-related protein required for olfaction. Genetics, 157(1), 211–224.

Coburn, C., & Gems, D. (2013). The mysterious case of the C. elegans gut granule: death fluorescence, anthranilic acid and the kynurenine pathway. Frontiers in genetics, 4, 151. doi:10.3389/fgene.2013.00151

Dan, K., Veetil, A. T., Chakraborty, K., & Krishnan, Y. (2019). DNA nanodevices map enzymatic activity in organelles. Nature Nanotechnology, 14(3), 252–259. doi:10.1038/s41565-019-0365-6

Dell’Angelica, E. C., Mullins, C., Caplan, S., & Bonifacino, J. S. (2000). Lysosome-related organelles. The FASEB Journal, 14(10), 1265–1278. doi:10.1096/fasebj.14.10.1265

Dittman, J., & Ryan, T. A. (2009). Molecular circuitry of endocytosis at nerve terminals. Annual Review of Cell and Developmental Biology, 25, 133–160. doi:10.1146/annurev.cellbio.042308.113302

Douglas, S. M., Bachelet, I., & Church, G. M. (2012). A logic-gated nanorobot for targeted transport of molecular payloads. Science, 335(6070), 831–834. doi:10.1126/science.1214081

Dumoulin, M., Conrath, K., Van Meirhaeghe, A., Meersman, F., Heremans, K., Frenken, L. G. J., … Matagne, A. (2002). Single-domain antibody fragments with high conformational stability. Protein Science, 11(3), 500–515. doi:10.1110/ps.34602

Ewert, S., Cambillau, C., Conrath, K., & Plückthun, A. (2002). Biophysical properties of camelid V(HH) domains compared to those of human V(H)3 domains. Biochemistry, 41(11), 3628–3636. doi:10.1021/bi011239a

Feng, Z., Li, W., Ward, A., Piggott, B. J., Larkspur, E. R., Sternberg, P. W., & Xu, X. Z. S. (2006). A C. elegans model of nicotine-dependent behavior: regulation by TRP-family channels. Cell, 127(3), 621–633. doi:10.1016/j.cell.2006.09.035

glr-4 (gene) - WormBase : Nematode Information Resource. (n.d.). Retrieved January 7, 2021, from https://wormbase.org/species/c_elegans/gene/WBGene00001615#0-9f31-10

Gottschalk, A., & Schafer, W. R. (2006). Visualization of integral and peripheral cell surface proteins in live Caenorhabditis elegans. Journal of Neuroscience Methods, 154(1-2), 68–79. doi:10.1016/j.jneumeth.2005.11.016

Granato, M., Schnabel, H., & Schnabel, R. (1994). pha-1, a selectable marker for gene transfer in C. elegans. Nucleic Acids Research, 22(9), 1762–1763. doi:10.1093/nar/22.9.1762

Haberland, M. E., & Fogelman, A. M. (1985). Scavenger receptor-mediated recognition of maleyl bovine plasma albumin and the demaleylated protein in human monocyte macrophages. Proceedings of the National Academy of Sciences of the United States of America, 82(9), 2693–2697.

Hamers-Casterman, C., Atarhouch, T., Muyldermans, S., Robinson, G., Hamers, C., Songa, E. B., … Hamers, R. (1993). Naturally occurring antibodies devoid of light chains. Nature, 363(6428), 446–448. doi:10.1038/363446a0

Hanson, P. I., Heuser, J. E., & Jahn, R. (1997). Neurotransmitter release - four years of SNARE complexes. Current Opinion in Neurobiology, 7(3), 310–315. doi:10.1016/S0959-4388(97)80057-8

Harmsen, M. M., & De Haard, H. J. (2007). Properties, production, and applications of camelid single-domain antibody fragments. Applied Microbiology and Biotechnology, 77(1), 13–22. doi:10.1007/s00253-007-1142-2

Hermann, G. J., Schroeder, L. K., Hieb, C. A., Kershner, A. M., Rabbitts, B. M., Fonarev, P., … Priess, J. R. (2005). Genetic analysis of lysosomal trafficking in Caenorhabditis elegans. Molecular Biology of the Cell, 16(7), 3273–3288. doi:10.1091/mbc.E05-01-0060

Hills, T., Brockie, P. J., & Maricq, A. V. (2004). Dopamine and glutamate control area-restricted search behavior in Caenorhabditis elegans. The Journal of Neuroscience, 24(5), 1217–1225. doi:10.1523/JNEUROSCI.1569-03.2004

Huizing, M., Helip-Wooley, A., Westbroek, W., Gunay-Aygun, M., & Gahl, W. A. (2008). Disorders of lysosome-related organelle biogenesis: clinical and molecular genetics. Annual Review of Genomics and Human Genetics, 9, 359–386. doi:10.1146/annurev.genom.9.081307.164303

Hulme, S. E., & Whitesides, G. M. (2011). Chemistry and the worm: Caenorhabditis elegans as a platform for integrating chemical and biological research. Angewandte Chemie, 50(21), 4774–4807. doi:10.1002/anie.201005461

Hunter, C. P., Winston, W. M., Molodowitch, C., Feinberg, E. H., Shih, J., Sutherlin, M., … Fitzgerald, M. C. (2006). Systemic RNAi in Caenorhabditis elegans. Cold Spring Harbor Symposia on Quantitative Biology, 71, 95–100. doi:10.1101/sqb.2006.71.060

Jani, M. S., Zou, J., Veetil, A. T., & Krishnan, Y. (2020). A DNA-based fluorescent probe maps NOS3 activity with subcellular spatial resolution. Nature Chemical Biology, 16(6), 660–666. doi:10.1038/s41589-020-0491-3

Jewett, J. C., Sletten, E. M., & Bertozzi, C. R. (2010). Rapid Cu-free click chemistry with readily synthesized biarylazacyclooctynones. Journal of the American Chemical Society, 132(11), 3688–3690. doi:10.1021/ja100014q

Jose, A. M., & Hunter, C. P. (2007). Transport of sequence-specific RNA interference information between cells. Annual Review of Genetics, 41, 305–330. doi:10.1146/annurev.genet.41.110306.130216

Kerk, S. Y., Kratsios, P., Hart, M., Mourao, R., & Hobert, O. (2017). Diversification of c. elegans motor neuron identity via selective effector gene repression. Neuron, 93(1), 80–98. doi:10.1016/j.neuron.2016.11.036

Kratsios, P., Pinan-Lucarré, B., Kerk, S. Y., Weinreb, A., Bessereau, J.-L., & Hobert, O. (2015). Transcriptional coordination of synaptogenesis and neurotransmitter signaling. Current Biology, 25(10), 1282–1295. doi:10.1016/j.cub.2015.03.028

Kratsios, P., Stolfi, A., Levine, M., & Hobert, O. (2011). Coordinated regulation of cholinergic motor neuron traits through a conserved terminal selector gene. Nature Neuroscience, 15(2), 205–214. doi:10.1038/nn.2989

Krishnan, Y., & Bathe, M. (2012). Designer nucleic acids to probe and program the cell. Trends in Cell Biology, 22(12), 624–633. doi:10.1016/j.tcb.2012.10.001

Krishnan, Y., Zou, J., & Jani, M. S. (2020). Quantitative imaging of biochemistry in situ and at the nanoscale. ACS central science, 6(11), 1938–1954. doi:10.1021/acscentsci.0c01076

Lee, H., Lytton-Jean, A. K. R., Chen, Y., Love, K. T., Park, A. I., Karagiannis, E. D., … Anderson, D. G. (2012). Molecularly self-assembled nucleic acid nanoparticles for targeted in vivo siRNA delivery. Nature Nanotechnology, 7(6), 389–393. doi:10.1038/nnano.2012.73

Li, J., Pei, H., Zhu, B., Liang, L., Wei, M., He, Y., … Fan, C. (2011). Self-assembled multivalent DNA nanostructures for noninvasive intracellular delivery of immunostimulatory CpG oligonucleotides. ACS Nano, 5(11), 8783–8789. doi:10.1021/nn202774x

Li, W., Koutmou, K. S., Leahy, D. J., & Li, M. (2015). Systemic RNA Interference Deficiency-1 (SID-1) Extracellular Domain Selectively Binds Long Double-stranded RNA and Is Required for RNA Transport by SID-1. The Journal of Biological Chemistry, 290(31), 18904–18913. doi:10.1074/jbc.M115.658864

Mahoney, T. R., Liu, Q., Itoh, T., Luo, S., Hadwiger, G., Vincent, R., … Nonet, M. L. (2006). Regulation of synaptic transmission by RAB-3 and RAB-27 in Caenorhabditis elegans. Molecular Biology of the Cell, 17(6), 2617–2625. doi:10.1091/mbc.E05-12-1170

Maxfield, F. R. (2014). Role of endosomes and lysosomes in human disease. Cold Spring Harbor Perspectives in Biology, 6(5), a016931. doi:10.1101/cshperspect.a016931

McEwan, D. L., Weisman, A. S., & Hunter, C. P. (2012). Uptake of extracellular double-stranded RNA by SID-2. Molecular Cell, 47(5), 746–754. doi:10.1016/j.molcel.2012.07.014

McGhee, J. D. (2007). The C. elegans intestine. Wormbook: the Online Review of C. Elegans Biology, 1–36. doi:10.1895/wormbook.1.133.1

Mellman, I., & Yarden, Y. (2013). Endocytosis and cancer. Cold Spring Harbor Perspectives in Biology, 5(12), a016949. doi:10.1101/cshperspect.a016949

Modi, S., M G, S., Goswami, D., Gupta, G. D., Mayor, S., & Krishnan, Y. (2009). A DNA nanomachine that maps spatial and temporal pH changes inside living cells. Nature Nanotechnology, 4(5), 325–330. doi:10.1038/nnano.2009.83

Modi, S., Nizak, C., Surana, S., Halder, S., & Krishnan, Y. (2013). Two DNA nanomachines map pH changes along intersecting endocytic pathways inside the same cell. Nature Nanotechnology, 8(6), 459–467. doi:10.1038/nnano.2013.92

Mukherjee, S., Ghosh, R. N., & Maxfield, F. R. (1997). Endocytosis. Physiological Reviews, 77(3), 759–803. doi:10.1152/physrev.1997.77.3.759

Murthy, K., Bhat, J. M., & Koushika, S. P. (2011). In vivo imaging of retrogradely transported synaptic vesicle proteins in Caenorhabditis elegans neurons. Traffic, 12(1), 89–101. doi:10.1111/j.1600-0854.2010.01127.x

Narayanaswamy, N., Chakraborty, K., Saminathan, A., Zeichner, E., Leung, K., Devany, J., & Krishnan, Y. (2019). A pH-correctable, DNA-based fluorescent reporter for organellar calcium. Nature Methods, 16(1), 95–102. doi:10.1038/s41592-018-0232-7

Nguyen, J. P., Shipley, F. B., Linder, A. N., Plummer, G. S., Liu, M., Setru, S. U., … Leifer, A. M. (2016). Whole-brain calcium imaging with cellular resolution in freely behaving Caenorhabditis elegans. Proceedings of the National Academy of Sciences of the United States of America, 113(8), E1074–81. doi:10.1073/pnas.1507110112

Nonet, M. L. (1999). Visualization of synaptic specializations in live C. elegans with synaptic vesicle protein-GFP fusions. Journal of Neuroscience Methods, 89(1), 33–40. doi:10.1016/s0165-0270(99)00031-x

Pereira, L., Kratsios, P., Serrano-Saiz, E., Sheftel, H., Mayo, A. E., Hall, D. H., … Hobert, O. (2015). A cellular and regulatory map of the cholinergic nervous system of C. elegans. eLife, 4. doi:10.7554/eLife.12432

Pérez, J. M., Renisio, J. G., Prompers, J. J., van Platerink, C. J., Cambillau, C., Darbon, H., & Frenken, L. G. (2001). Thermal unfolding of a llama antibody fragment: a two-state reversible process. Biochemistry, 40(1), 74–83. doi:10.1021/bi0009082

Rizzoli, S. O. (2014). Synaptic vesicle recycling: steps and principles. The EMBO Journal, 33(8), 788–822. doi:10.1002/embj.201386357

Rothaug, M., Zunke, F., Mazzulli, J. R., Schweizer, M., Altmeppen, H., Lüllmann-Rauch, R., … Blanz, J. (2014). LIMP-2 expression is critical for β-glucocerebrosidase activity and α-synuclein clearance. Proceedings of the National Academy of Sciences of the United States of America, 111(43), 15573–15578. doi:10.1073/pnas.1405700111

Saha, S., Prakash, V., Halder, S., Chakraborty, K., & Krishnan, Y. (2015). A pH-independent DNA nanodevice for quantifying chloride transport in organelles of living cells. Nature Nanotechnology, 10(7), 645–651. doi:10.1038/nnano.2015.130

Saminathan, A., Devany, J., Veetil, A. T., Suresh, B., Pillai, K. S., Schwake, M., & Krishnan, Y. (2020). A DNA-based voltmeter for organelles. Nature Nanotechnology. doi:10.1038/s41565-020-00784-1

Schroeder, L. K., Kremer, S., Kramer, M. J., Currie, E., Kwan, E., Watts, J. L., … Hermann, G. J. (2007). Function of the Caenorhabditis elegans ABC transporter PGP-2 in the biogenesis of a lysosome-related fat storage organelle. Molecular Biology of the Cell, 18(3), 995–1008. doi:10.1091/mbc.e06-08-0685

Seeman, N. C., & Sleiman, H. F. (2017). DNA nanotechnology. Nature Reviews Materials, 3(1), 17068. doi:10.1038/natrevmats.2017.68

Sharma, S., Zaveri, A., Visweswariah, S. S., & Krishnan, Y. (2014). A fluorescent nucleic acid nanodevice quantitatively images elevated cyclic adenosine monophosphate in membrane-bound compartments. Small (Germany), 10(21), 4276–4280. doi:10.1002/smll.201400833

Soukas, A. A., Carr, C. E., & Ruvkun, G. (2013). Genetic regulation of Caenorhabditis elegans lysosome related organelle function. PLoS Genetics, 9(10), e1003908. doi:10.1371/journal.pgen.1003908

Stefanakis, N., Carrera, I., & Hobert, O. (2015). Regulatory Logic of Pan-Neuronal Gene Expression in C. elegans. Neuron, 87(4), 733–750. doi:10.1016/j.neuron.2015.07.031

Südhof, T. C. (1995). The synaptic vesicle cycle: a cascade of protein-protein interactions. Nature, 375(6533), 645–653. doi:10.1038/375645a0

Surana, S., Bhat, J. M., Koushika, S. P., & Krishnan, Y. (2011). An autonomous DNA nanomachine maps spatiotemporal pH changes in a multicellular living organism. Nature Communications, 2, 340. doi:10.1038/ncomms1340

Teschendorf, D., & Link, C. D. (2009). What have worm models told us about the mechanisms of neuronal dysfunction in human neurodegenerative diseases? Molecular Neurodegeneration, 4, 38. doi:10.1186/1750-1326-4-38

Thekkan, S., Jani, M. S., Cui, C., Dan, K., Zhou, G., Becker, L., & Krishnan, Y. (2019). A DNA-based fluorescent reporter maps HOCl production in the maturing phagosome. Nature Chemical Biology, 15(12), 1165–1172. doi:10.1038/s41589-018-0176-3

van der Linden, R. H., Frenken, L. G., de Geus, B., Harmsen, M. M., Ruuls, R. C., Stok, W., … Verrips, C. T. (1999). Comparison of physical chemical properties of llama VHH antibody fragments and mouse monoclonal antibodies. Biochimica et Biophysica Acta, 1431(1), 37–46. doi:10.1016/s0167-4838(99)00030-8

Veetil, A. T., Chakraborty, K., Xiao, K., Minter, M. R., Sisodia, S. S., & Krishnan, Y. (2017). Cell-targetable DNA nanocapsules for spatiotemporal release of caged bioactive small molecules. Nature Nanotechnology, 12(12), 1183–1189. doi:10.1038/nnano.2017.159

Veetil, A. T., Zou, J., Henderson, K. W., Jani, M. S., Shaik, S. M., Sisodia, S. S., … Krishnan, Y. (2020). DNA-based fluorescent probes of NOS2 activity in live brains. Proceedings of the National Academy of Sciences of the United States of America, 117(26), 14694–14702. doi:10.1073/pnas.2003034117

Winston, W M, Molodowitch, C., & Hunter, C. P. (2002). Systemic RNAi in *C. elegans* requires the putative transmembrane protein SID-1. Science, 295(5564), 2456–2459. doi:10.1126/science.1068836

Winston, William M, Sutherlin, M., Wright, A. J., Feinberg, E. H., & Hunter, C. P. (2007). Caenorhabditis elegans SID-2 is required for environmental RNA interference. Proceedings of the National Academy of Sciences of the United States of America, 104(25), 10565–10570. doi:10.1073/pnas.0611282104

Zhao, Y.-X., Shaw, A., Zeng, X., Benson, E., Nyström, A. M., & Högberg, B. (2012). DNA origami delivery system for cancer therapy with tunable release properties. ACS Nano, 6(10), 8684–8691. doi:10.1021/nn3022662

## Bibliography

1. Paredes, E. & Das, S. R. Click chemistry for rapid labeling and ligation of RNA. Chembiochem 12, 125–131 (2011).

2. Jewett, J. C., Sletten, E. M. & Bertozzi, C. R. Rapid Cu-free click chemistry with readily synthesized biarylazacyclooctynones. J. Am. Chem. Soc. 132, 3688–3690 (2010).

3. Baskin, J. M. et al. Copper-free click chemistry for dynamic in vivo imaging. Proc. Natl. Acad. Sci. USA 104, 16793–16797 (2007).

4. Milligan, J. F., Groebe, D. R., Witherell, G. W. & Uhlenbeck, O. C. Oligoribonucleotide synthesis using T7 RNA polymerase and synthetic DNA templates. Nucleic Acids Res. 15, 8783–8798 (1987).

5. Huang, F., He, J., Zhang, Y. & Guo, Y. Synthesis of biotin-AMP conjugate for 5’ biotin labeling of RNA through one-step in vitro transcription. Nat. Protoc. 3, 1848–1861 (2008).

6. Merritt, C., Rasoloson, D., Ko, D. & Seydoux, G. 3’ UTRs are the primary regulators of gene expression in the C. elegans germline. Curr. Biol. 18, 1476–1482 (2008).

7. Brenner, S. Caenorhabdztzs elegans. 71–94 (1974).

8. Mahoney, T. R. et al. Regulation of synaptic transmission by RAB-3 and RAB-27 in Caenorhabditis elegans. Mol. Biol. Cell 17, 2617–2625 (2006).

9. Bounoutas, A., Zheng, Q., Nonet, M. L. & Chalfie, M. mec-15 encodes an F-box protein required for touch receptor neuron mechanosensation, synapse formation and development. Genetics 183, 607–17, 1SI (2009).

10. Nguyen, J. P. et al. Whole-brain calcium imaging with cellular resolution in freely behaving Caenorhabditis elegans. Proc. Natl. Acad. Sci. USA 113, E1074–81 (2016).

11. Altun-Gultekin, Z. et al. A regulatory cascade of three homeobox genes, ceh-10, ttx-3 and ceh-23, controls cell fate specification of a defined interneuron class in C. elegans. Development 128, 1951–1969 (2001).

12. Treusch, S. et al. Caenorhabditis elegans functional orthologue of human protein h-mucolipin-1 is required for lysosome biogenesis. Proc. Natl. Acad. Sci. USA 101, 4483–4488 (2004).

13. Zhang, S. O., Trimble, R., Guo, F. & Mak, H. Y. Lipid droplets as ubiquitous fat storage organelles in C. elegans. BMC Cell Biol. 11, 96 (2010).

14. Mello, C. C., Kramer, J. M., Stinchcomb, D. & Ambros, V. Efficient maintenance. 10, 3959–3970 (1991).

15. Granato, M., Schnabel, H. & Schnabel, R. *pha-1*, a selectable marker for gene transfer in *C.elegans*. Nucleic Acids Res. 22, 1762–1763 (1994).

16. Stinchcomb, D. T., Shaw, J. E., Carr, S. H. & Hirsh, D. Extrachromosomal DNA transformation of Caenorhabditis elegans. Mol. Cell. Biol. 5, 3484–3496 (1985).

17. Single worm PCR | Schedl Lab. at <http://genetics.wustl.edu/tslab/protocols/genomic-stuff/single-worm-pcr/>

18. Sauvola, J. & Pietikäinen, M. Adaptive document image binarization. Pattern Recognit 33, 225–236 (2000).

